# *k*-mer-based GWAS in a wheat collection reveals novel and diverse sources of powdery mildew resistance

**DOI:** 10.1101/2024.10.03.616421

**Authors:** Benjamin Jaegle, Yoav Voicheck, Max Haupt, Alexandros G. Sotiropoulos, Kevin Gauthier, Matthias Heuberger, Esther Jung, Gerhard Herren, Victoria Widrig, Rebecca Leber, Yipu Li, Beate Schierscher, Sarah Serex, Maja Boczkowska, Marta-Puchta Jasińska, Paulina Bolc, Boulos Chalhoub, Nils Stein, Beat Keller, Javier Sanchez Martin

## Abstract

**Background:** Wheat landraces and cultivars stored in gene banks worldwide represent a valuable source of genetic diversity for discovering genes critical for agriculture, which is increasingly constrained by climate change and inputs reduction. We assembled and genotyped, using DArTseq technology, a panel of 461 accessions representative of the genetic diversity of Swiss wheat material. The collection was evaluated for powdery mildew resistance under field conditions for two consecutive years and at the seedling stage with 10 different wheat powdery mildew isolates.

**Results:** To identify the genetic basis of mildew resistance in wheat, we developed a *k*-mer-based GWAS approach using multiple fully-assembled genomes including *Triticum aestivum* as well as four progenitor genomes. Compared to approaches based on single reference genomes, we unambiguously mapped an additional 25% resistance-associated *k*-mers. Our approach outperformed SNP-based GWAS in terms of number of loci identified and precision of mapping. In total, we detected 34 (*Pm*) powdery mildew resistance loci, including seven previously-described and more importantly 27 novel loci active at the seedling stage. Furthermore, we identified a region associated with adult plant resistance, which was not detected with SNP-based approaches.

**Conclusions:** The described non-reference-based approach highlights the potential of integrating multiple wheat reference genomes with *k*-mer GWAS to harness the untapped genetic diversity present in germplasm collections.

## Introduction

Wheat production, which accounts for 18% of global calories intake, is reduced by 20% annually due to pests and diseases [1]. One significant threat is wheat powdery mildew, caused by the obligate biotrophic ascomycete *Blumeria graminis* f. sp*. tritici*. This pathogen can reduce grain yield by 7.6-19.9% [2], leading to annual losses exceeding 4 billion euros worldwide despite the use of agrochemicals, adaptation of agronomic practices and the deployment of resistance cultivars. Although more than a hundred powdery mildew resistance genes (*Pm*) have been reported, only a few can provide effective resistance in the main wheat-growing areas. For example, from the 16 molecularly cloned *Pm* genes, only a handful are efficient against wheat mildew races in three of the 17 Agroecological Zones where wheat is grown, leaving vast regions, like Europe and Central Asia with no effective *Pm* resistance genes [3], forcing breeders to rely on challenging massive evaluation of overall resistance, which remain challenging to implement in breeding programmes due to the underlying genetic complexity.

This moderate efficacy of powdery mildew resistance loci in modern varieties is the result of pathogen adaptation as well as a consequence of the narrow genetic base imposed by the bottlenecks of hexaploidization and domestication, exacerbated during the Green Revolution with breeding activities based on few founders. This resulted in genetic erosion and increasing susceptibility and vulnerability to environmental stresses, pests, and diseases, forcing farmers to increasingly rely on pesticides to control wheat diseases, such as powdery mildew. However, chemical control via pesticides is costly, harmful to ecosystems, and increasingly ineffective due to global fungicide resistance [4]. Additionally, the European Commission aims to reduce pesticide usage by 50% by 2030 [5]. In this context, resistance breeding is critical for sustainably controlling pests and pathogens while reducing pesticide dependency.

Plant disease resistance is molecularly diverse and categorized as either race-specific or quantitative resistance (QR). Race-specific resistance provides mostly complete resistance to some pathogen races and only occurs in the presence of a resistance (*R*) gene in the plant, and the corresponding effector-encoding avirulence (*Avr*) gene in the pathogen [6]. In contrast, QR provides partial quantitative resistance at the adult plant stage to all races of a pathogen species, independent of rapidly evolving pathogen effectors [7]. There are only ∼ 460 wheat resistance genes genetically defined [8] that are currently being used in breeding. However, based on pan-genome analyses, not representative of the all wheat diversity, up to 7,000 resistance genes could be present in the wheat gene pool (Walkowiak et al., 2020). This means that there is a putatively large, unexplored diversity of *Pm* genes stored in wheat gene banks that await being uncovered and used for resistance breeding. In this context, crop wild relatives (CWR) and landraces, genetically diverse populations traditionally cultivated in low-input systems and adapted to different ecoclimatic conditions, represent a valuable genetic resource for improving crops against the occurring climate change.

As an alternative to traditional quantitative trait locus (QTL) mapping, genome-wide association studies (GWAS) offer a faster approach to identifying statistical associations between phenotypic and genetic variations. This method bypasses the need to create segregating populations, significantly reducing the time required. However, there is a notable lack of assessment of the genetic and, in particular, phenotypic diversity of CWR and landraces, limiting their use in breeding [9]. Although genome sequencing costs have dramatically declined, generating reference-quality assemblies of a species like wheat, with its 15Gb genome, remains both expensive and computationally demanding. Alternatively, various array- and sequencing-based genotyping platforms have emerged to interrogate the genome-wide diversity of wheat genomes. One such method is single-nucleotide polymorphism (SNP)-based arrays, which have proven useful in identifying disease resistance genes in wheat germplasms. However, they typically rely on a single reference genome (or a few accessions), dramatically limiting the SNP set that can be detected [10], and they only can detect SNPs but not the remaining structural variations (SVs), such as insertions, deletions, duplications, copy number variants (CNVs) or translocations that have been shown to underlie relevant traits, such as stress tolerance or disease resistance [11].

To overcome these limitations inherent to SNP-based arrays, alternative genotyping approaches were developed, e.g. Diversity Array Technology sequencing, or DArTseq (http://www.diversityarrays.com/), which produces short sequence fragments by restriction enzyme-mediated genome complexity reduction [12]. This technology selects predominantly low-copy number regions of the genome and by sequencing small regions, typically 200-300 bp, with high coverage, it enables the calling of high-quality SNPs [13]. The resulting SNPs information allows interrogating the diversity of hundreds of wheat genotypes at an affordable cost. Such an approach has been successfully applied to wheat in multiple studies [9,14].

In many model species, only one reference genome is used, and wheat is no exception. Most studies involving wheat genotyping define SNPs or other variations relative to the *Chinese Spring* (CS) reference genome. GWAS based on single reference genomes is restricted to the discovery of genes and variations present in that specific reference genome [15]. However, plants have been shown to have extensive structural variation [16,17] which cannot be represented by a single reference genome, highlighting the need for more diversity. Further, resistance genes are often part of introgressions from wild relatives or they show presence/absence polymorphism. Consequently, a single reference genome is unlikely to capture all genetic variants [18,19]. To avoid these two biases one can use multiple reference genomes to increase the variation detected [20].

Alternatively, an alignment-free approach can be used. This approach differs from the classical SNPs-based markers as it directly correlates the presence/absence of small sequences (typically 31bp), called *k*-mers, with phenotypes. Using *k*-mers as markers allows the detection of almost any type of structural variant, including insertions, deletions, or transpositions in addition to classical SNPs [21]. However, one limitation of alignment-free approaches is the challenge of linking *k*-mers with causal genes.

To leverage the untapped genetic diversity of wheat accessions and establish a broadly applicable workflow for plant genomics (**Figure 1**), we assembled a diverse collection of 461 Swiss bread wheat accessions (**Figure 1A**) and developed a *k*-mer-based GWAS pipeline using multiple wheat reference genomes inspired by work done in flowering plants [21] (**Figure 1B-D**). We mapped the raw reads onto ten *T. aestivum* reference genomes generated in the 10+ Wheat Genome Project [22], as well as multiple wheat progenitors (**Figure 1E**). With this approach, we demonstrated that we could detect a larger diversity of segregating loci involved in powdery mildew resistance that would have been missed using a single reference genome. The association mapping detected multiple known *Pm* genes including *Pm1*, *Pm2*, *Pm60*, or *Pm4b*, but also novel regions associated with powdery mildew resistance in chromosomes 3D, 5D, and 6A, totaling 34 potential genomic regions of interest spread across all the subgenomes.

**Figure 1:**
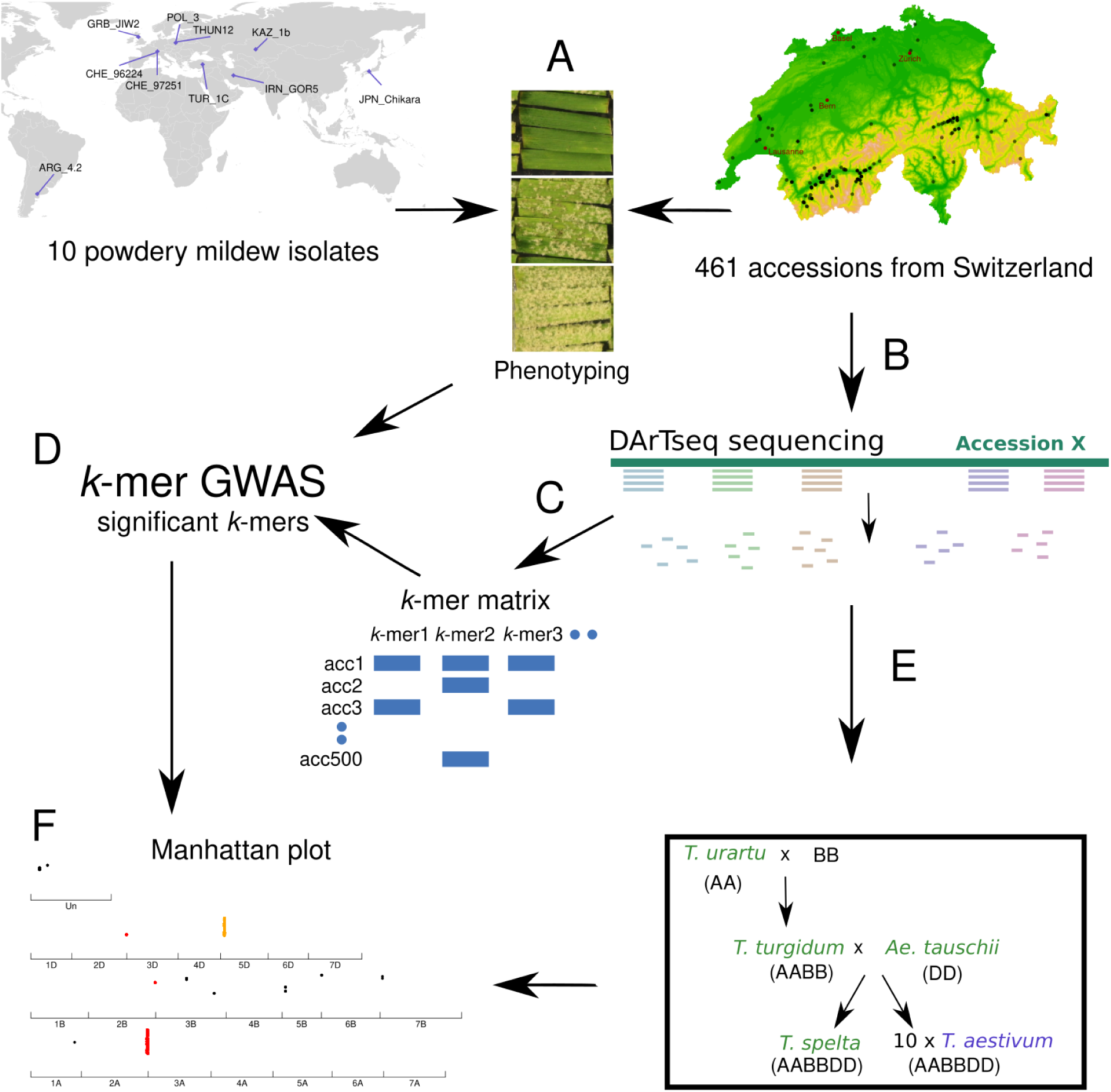
Workflow to identify the genetic basis of resistance in the wheat-powdery mildew pathosystem. A: All accessions from the Swiss wheat collection were phenotyped using 10 *Bgt* isolates from around the globe. B: All accessions from the collection were sequenced using DArTseq. C: From the raw reads of the DArTseq, 31bp *k*-mers were generated, and a presence/absence matrix was used to run GWAS. D: Using the *k*-mer matrix and the phenotyping data, GWAS was used to find *k*-mers significantly associated with the phenotype. E: All the DArTseq raw data were mapped to ten *Triticum aestivum* genomes as well as three progenitor genomes and the genome of *T. spelta*. F: Manhattan plots were generated for each genome of reference. The significant peaks were extracted to select candidate genes.

Of note, unlike standard GWAS, our method identifies associations with structural variations and sites not present in a single reference genome, highlighting the relevance of landraces and old cultivars stored in genebanks as a source of novel genetic variation important not only for mildew resistance but any other agronomic trait important for adaptation to a changing environment.

## Results

### A Swiss wheat collection shows variation in powdery mildew resistance to 10 *Bgt* **isolates with different virulence profiles**

Powdery mildew isolates differ in avirulence gene content and, therefore, the observed reaction to wheat *Pm* resistance genes is isolate-dependent. High-throughput sequencing technologies allow a prediction of effector gene content. As most avirulence genes are still unknown at the molecular level, there are no straightforward tools to select *Bgt* isolates with functional effectors without prior functional validation or extensive phenotyping in tester lines. Therefore, we picked *Bgt* isolates from different regions where wheat is cultivated to support a wide search of potential host-resistance components (**Figure 2A**).

**Figure 2:**
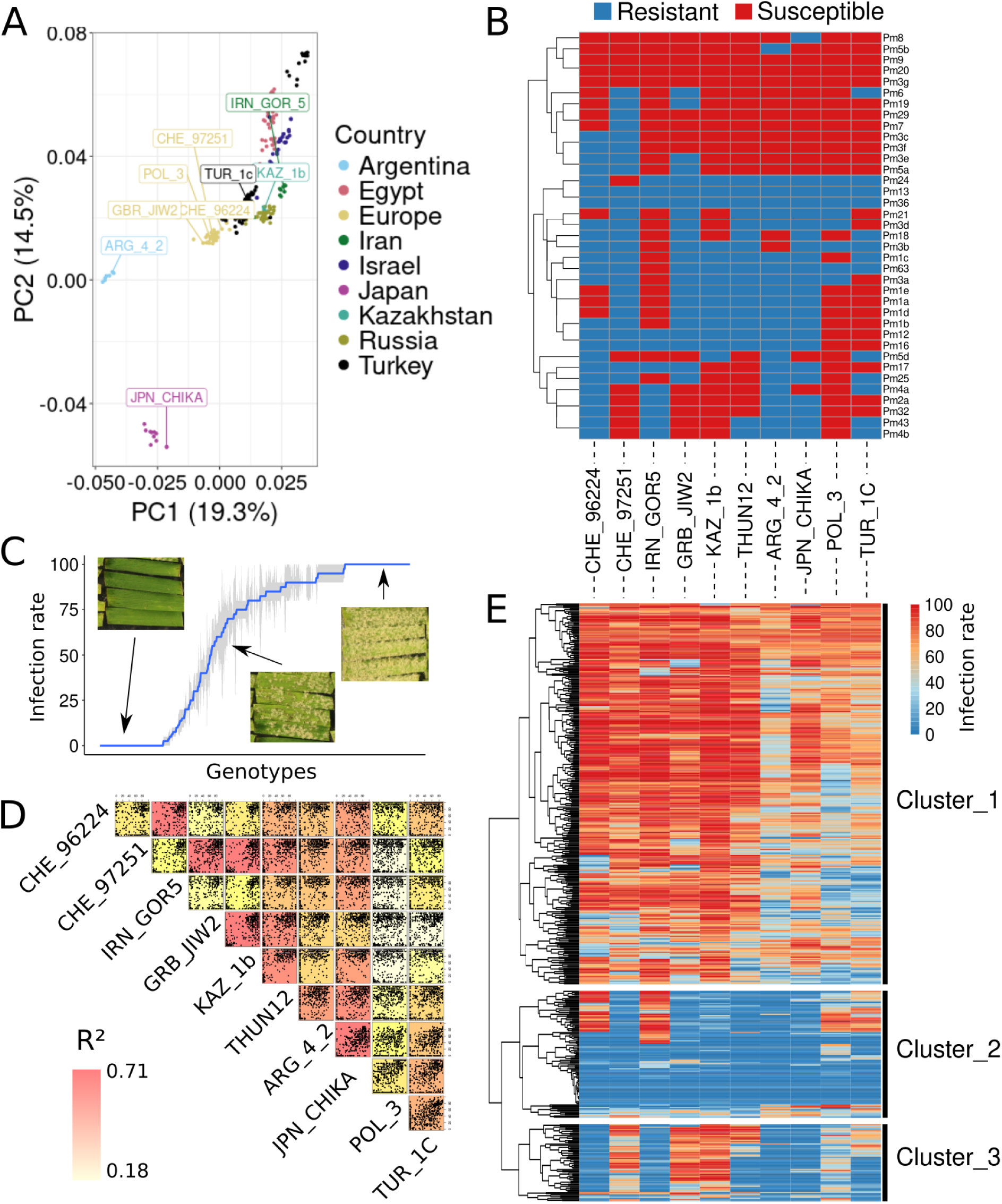
Phenotyping of the Swiss wheat collection with 10 powdery mildew isolates representative of the global genetic diversity of *Blumeria graminis* f. sp. *tritici*. **A**: PCA of 400 *Bgt* isolates with 9 of the 10 isolates used in this study highlighted. **B**: Avirulence (blue) and virulence (red) pattern of the 10 isolates across 37 Pm-tester lines [89]. **C:** Phenotype distribution of the isolate CHE_96224 on the Swiss wheat collection. Pictures represent example phenotypes for fully resistant, partially resistant, and fully susceptible seedling reactions. **D**: Correlation plot of the phenotype of all accessions for each isolate. Background color represents the Pearson correlation value. E: Heatmap representing the phenotype of each accession of the Swiss collection for the ten *Bgt* isolates sorted the same way as B. The three main clusters were split.

We selected nine *Blumeria graminis* f. sp. *tritici* isolates from six of the nine populations representing geographical origins detected by [23]. These include two Swiss isolates (CHE_96224 and CHE_97251) from the European cluster, and one from England, (GRB_JIW2) Poland (POL_3), Turkey (TUR_1c), Iran (IRN_GOR5), Kazakhstan (KAZ_1b), Argentina (ARG_4_2), and Japan (JPN_CHIKA) each (**Figure 2A**). An additional isolate from Poland (THUN12), is a hybrid between wheat and rye mildew, which was not part of the PCA analysis presented in Fig. 2A [23]. The geographic origin of those 10 isolates is shown in **Figure 1A**.

To assess the potential of the selected *Bgt* isolates to search for resistance genes, we first determined their avirulence/virulence pattern on 37 *Pm*-tester lines (**Figure 2B)**. Such tester genotypes have been generated through different crosses, usually involving a wild relative of wheat with a susceptible hexaploid wheat genotype, to ensure the sole presence of a specific *Pm* gene for pathogenicity tests. Across all *Pm*-tester lines, each of the ten isolates revealed a different resistance pattern. Interestingly, only the *Pm13* tester line was resistant to all isolates. In contrast, *Pm9*, *Pm5b*, *Pm20*, and *Pm3g* were susceptible to all ten isolates. Overall, we observed a large variation between the *Pm*-tester lines across the isolates, demonstrating the potential of the chosen *Bgt* isolates for detecting a wide variety of powdery mildew resistance genes (**Figure 2B**).

Using the ten chosen *Bgt* isolates, we then phenotyped a collection of Swiss accessions (**Supplemental Table**), consisting of 139 and 276 spring and winter accessions, respectively, together with 46 accessions of undefined growth habit (growing in both conditions), totalling 461 accessions. Those accessions include old cultivars and landraces and represent a selection of the diversity held in the Swiss genebank. Mildew resistance phenotyping revealed a large variation in powdery mildew resistance among the collection (**Figure 2E**). Across the 10 isolates, an average of 80 accessions were resistant to individual *Bgt* isolates, while, on average, 354 were susceptible to single *Bgt* isolates (**Figure 2E**). The most virulent *Bgt* isolate, POL_3, had only 32 resistant accessions, while the most avirulent *Bgt* isolate, ARG_4_2, had over 25% (n = 146) of the accessions showing resistance. Additionally, 206 accessions were susceptible to all *Bgt* isolates, while only 28 were resistant to all. The latter are of great interest to discover potential broad-spectrum resistance genes.

To further explore the commonality between the isolates, we looked at the correlation of their virulence/avirulence patterns across the Swiss collection. (**Figure 2D**). The highest correlations were observed between JPN_CHIKA and ARG_4_2 (R=0.76) and between CHE_96224 and IRN_GOR5 (R²=0.81), with POL_3 being the most distinct *Bgt* isolate with an average correlation value of 0.42 to all other isolates. There was no particular correlation between the geographical origin of the isolate and the virulence/avirulence pattern. For example, the virulence/avirulence patterns of the two isolates from Switzerland, as well as the two from Poland, are poorly correlated with each other compared to the other isolates. When comparing the genetic distance of powdery mildew isolates, there was no clear correlation between the phenotypic and genotypic variation **(Figure 2A, D).**

We developed a Shiny app (https://benjiapp.shinyapps.io/Map_agent_pheno/) to explore the resistance distribution for each *Bgt* isolate. As an example, the resistance distribution is shown for isolate CHE_96224, a very avirulent *Bgt* isolate widely used [24,25] (**Figure 2C).** 138 accessions showed resistance (<=20% leaf covered) to CHE_96224, whereas 94 accessions were fully susceptible (=100), with 229 accessions showing an intermediate phenotype.

When grouping based on the phenotypic responses to *Bgt* isolates, wheat accessions were grouped into three main clusters. The first and biggest cluster (315 accessions), consists of mostly susceptible accessions to all *Bgt* (**Figure 2E**). The second cluster (105 accessions), has mostly accessions resistant to all isolates or accessions susceptible to CHE_96224, IRN_GOR5, POL_3, TUR_1C *Bgt* isolates. When comparing with the *Pm*-tester lines, such a pattern of accessions being only susceptible to CHE_96224, IRN_GOR5, POL_3, and TUR_1C perfectly matches with the virulence/avirulence patterns of the lines containing *Pm1e*, *a*, and *d,* which suggests that these accessions might contain the *Pm1* locus. The third cluster was formed by 64 accessions that are all resistant to CHE_96224, IRN_GOR5, ARG_4_2, and JPN_CHIKA, while a subset of them are also resistant to THUN12 and TUR_1C. Comparison with the *Pm*-tester lines resistance spectra shows that such a pattern corresponds to *Pm4b* and *Pm2* or *Pm32*-containing lines and indicates that the *Pm2*, *Pm4b*, or *Pm32* genes are likely to be present in the Swiss collection.

### The *Pm2* and *Pm4b* race-specific resistance genes are widely present in the Swiss collection

*Pm4* and *Pm2* are key genes used in breeding programs involved in powdery mildew resistance [26,27] and have been detected in several wheat populations globally [28,29]. To investigate whether local wheat hexaploid landraces and old cultivars from the Swiss collection contain these genes as suggested by the phenotyping analysis described above, we tested for the presence of *Pm4b* and *Pm2* using haplotype markers and further sequencing [26,30]. Out of 461 accessions, 50 and 31 contained *Pm2* and *Pm4b*, respectively, with eight accessions having both genes. Whereas most of the accessions containing *Pm4b* showed the expected resistance to CHE_96224, IRN_GOR5, JPN_CHIKA, THUN12, and TUR_1C [26] few accessions did not match this pattern (**Supplemental Figure 1**). In the case of expanded resistance, such accessions might contain additional resistance genes, while the lack of resistance is most likely due to the presence of the non-functional *Pm4f* allele [26,31].

On the other hand, most accessions containing the *Pm2* gene had a resistance spectrum matching the expected pattern of the *Pm2* near isogenic lines (NIL), being resistant against CHE_96224, IRN_GOR5, ARG_4_2, and JPN_CHIKA. Some exceptions were observed, which are possibly caused by the presence of other resistance genes. Given these two examples, it is very likely that additional sources of resistance are present in the Swiss wheat collection. To discover new genes involved in powdery mildew resistance we set out to run GWAS analysis.

### DArTseq genotyping reveals large genetic diversity of the Swiss collection

All accessions from the Swiss wheat collection were genotyped using DArTseq. This genotyping by sequencing (GBS) method aims to reduce the proportion of repetitive sequences, keeping genome coverage more homogenous [12]. To assess this, we analyzed the distribution of the sequencing reads across the Chinese Spring IWGSC_2.1 reference genome (CS, [32]). Mapping raw reads of all accessions to the reference genome revealed that 40% of the annotated genes had at least one read within their coding region. When extending to +/- 10 kb around the coding sequence of a gene, 72% of the genes were tagged (**Supplemental Figure 2**). Additionally, the distribution of reads was similar between homologous chromosomes with 1,491,774, 1,605,237, and 1,350,862 reads mapping to the subgenomes A, B, and D, respectively. **(Supplemental Figure 3)**. On average, 2.3% of the reads per accession did not map to the CS genome. AG-392 had the fewest mapped reads, with 8.8 % not mapping to the CS genome.

The collection was also genotyped with the 12k Illumina Infinium 15K wheat SNP array (TraitGenetics GmbH, Gatersleben, Germany). After filtering, 11,252 SNPs were kept (see Methods). In addition to the SNP array, a matrix of 10,068 SNPs was generated by mapping the DArTseq reads to the CS genome. When comparing the number of SNPs per chromosome for each matrix, the D subgenome showed the expected lower number of SNPs **(Supplemental Figure 4)**. This reduced diversity is due to a strong bottleneck in the formation of hexaploid wheat [33]. In addition to the two SNPs matrices, we generated a presence/absence matrix of 176 million unique *k*-mers of length 31bp. We then used these two SNP-based matrices (DArTseq, the SNP-chip,) and the *k*-mers matrix - to perform GWAS.

### *K*-mer-based association mapping outperforms GWAS based on SNP matrices

To perform a comparative analysis of the performance of GWAS when using the different genotyping methods, we set up a pipeline allowing the use of the three matrices and the mildew resistance as phenotype. The GWAS results using the two SNP matrices showed very similar results **(Supplemental Figures 5 and 6**). With both SNPs matrices, for the *Bgt* isolate POL_3, no significant regions were detected. However, combining the GWAS results from all the other isolates, we detected 241 and 53 significant SNPs from the DArTseq and the SNP-chip, respectively. They were embedded in seven genomic regions, with two regions standing out above the rest: one located at the beginning of chromosome 5D for resistance to CHE_96224 and IRN_GOR5, and one at the end of chromosome 7A for resistance to ARG_4_2, CHE_97251, GRB_JIW2, JPN_CHIKA, KAZ_1b, and THUN12. **(Figure 3A)**. Those two regions are known to contain *Pm* genes: *Pm2*, located at the beginning of chromosome 5D around 43.4 Mb [34], and *Pm1* and *Pm60* are located at the end of chromosome 7A [35,36]. We observed that the corresponding regions on homoeologous chromosomes also showed significantly associated SNPs. This is most likely due to the homology of sequences between the chromosomes and/or linkage disequilibrium. Based on the phenotyping clustering as well as the comparison with *Pm* tester lines, *Pm1/Pm60* and *Pm2* were expected to be present in the Swiss wheat collection. Surprisingly, we did not detect *Pm4* using the SNPs matrices.

**Figure 3:**
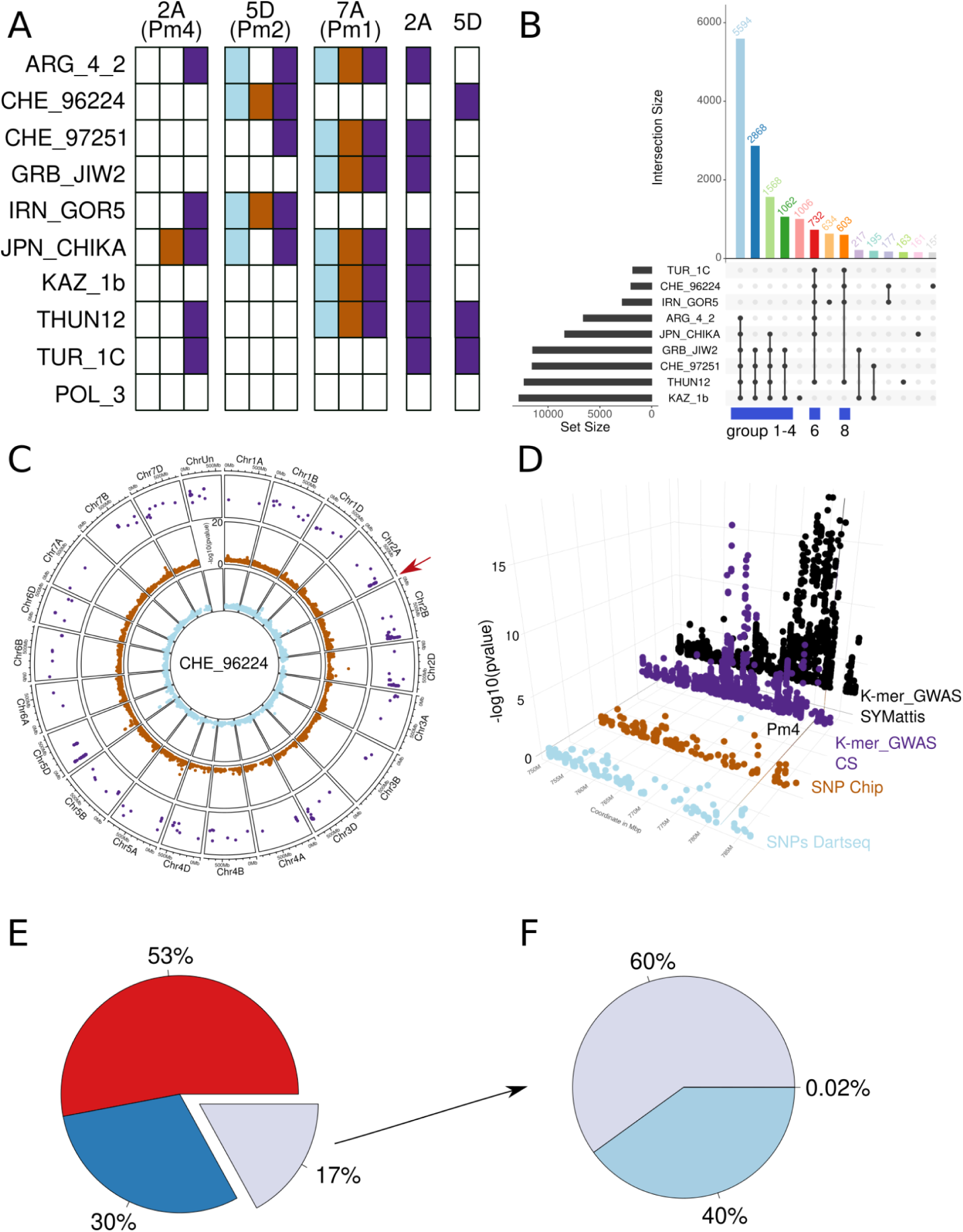
*k*-mer GWAS results. **A:** Summary of the GWAS results based on CS for *Pm1*, *Pm2* and *Pm4* for all three matrices for all the isolates, as well as two regions only found using *k*-mers. The colors blue, brown and purple respectively represent the SNP-Chip, the SNPs generated from the DArTseq data and the *k-*mers. Filled squares represent the presence of an associated region. **B**: UpSet plot for all the significant *k*-mers of each of the 10 isolates. The biggest set of common significant *k*-mers (5,594) was found between the isolates ARG_4_2 and JPN_CHIKA, GRB_JIW2, CHE_97251, THUN12, and KAZ_1b. The first 14 groups are colored. The entire data set can be explored in the shiny app at https://benjiapp.shinyapps.io/Manhattan_plot/. **C**: Circular plot representing Manhattan plots comparing three matrices used for the GWAS of *Bgt* isolate CHE_96224. The inner circle in blue represents the SNP matrix generated using DArTseq, and the middle circle in orange represents the matrix from the SNP chip. The outer circle represents the *k*-mer-GWAS where only significant *k*-mers are displayed. The y-axis is the same for all three plots. **D**: 3D dot plot of the different Manhattan plots for the region of *Pm4* (red arrow on C). The color legends are as in A. The *k*-mer-GWAS SYMattis represent the *k*-mer-GWAS using the SYMattis genomes as a reference. The line is at the position of the *Pm4* gene in the SYMattis genome. **E:** Proportion of *k*-mers mapping to all the *Triticum aestivum* genomes (red), some of the genomes (blue), and none of the genomes (grey). **F**: Proportion of reads that do not map to any of the *T. aestivum* genomes, but map to genomes of wheat progenitors or relatives. The other 40% do not map to any (light blue) and only 0.02% map to all four genomes.

To test whether the output of the GWAS analysis could be improved by using a reference-free approach, we adapted a previously published *k*-mer GWAS approach [21]. The GWAS using *k*-mers as genotype yielded 16,895 significant *k*-mers with an average of 5,876 significant *k*-mers per isolate. Isolate KAZ_1b had the highest number of significant *k*-mers (12,806), and TUR_1C lowest number (1,819) (**Figure 3B**). Similar to the SNPs, no significant *k*-mers were detected for the POL_3 isolate (summary in **Supplemental Table**). Using the SNPs matrices, the classical outputs are lists of significant SNPs, for *k*-mers, the output is a list of significant *k*-mers, of 31 bp in length. To position these *k*-mers within a genome, several steps are needed: after retrieving the reads containing the *k*-mers, these reads are aligned to the genome and then the k-mers p-value can be linked to a genome position (see **Figure 1** for the pipeline).

*k*-mer GWAS identified the same resistance-associated regions as the two SNPs matrices, plus additional novel associations. For example, we detected an associated region on chromosome 2A that corresponds to the location of the *Pm4* gene (**Figure 3C,D**). Based on the resistance pattern analysis of the phenotypic data and the haplotype analysis (**Supplemental Figure 1**), this gene is present in the Swiss wheat collection. To better understand why such a region is only detected using *k*-mers, we zoomed into the *Pm4* region and plotted the results of the three approaches together (**Figure 3D**). This revealed that the higher resolution of the *k*-mer GWAS was due to the higher number of markers in the region: there are 165, 141, and 1627 markers for the DArTseq, the SNP-chip, and the *k*-mer-GWAS, respectively.

Only with the *k*-mers GWAS approach, we detected additional associated regions in chromosomes 2A and 5D (**Figure 3A).** In total, we found 21 associated regions that do not overlap with known *Pm* genes. The Manhattan plot containing all the significantly associated regions for all the isolates is presented in **Supplemental Figure 7.** These results demonstrate that *k*-mer-based GWAS outperforms the approaches based on SNPs, and the comparative analysis highlights that the increased number of markers is mostly the reason for this improvement.

### Multiple reference genomes improve the accuracy of significant *k*-mers positions and refine genomic regions definitions

Of the 16,895 significant k-mers identified, 68% (11,570) were mapped to the CS reference genome. In contrast, while only 2.3% of the overall reads did not map to the CS genome, a much larger proportion, i.e.32% of the significant *k*-mers, did not map to the CS genome. This suggests a significant enrichment of non-mapping k-mers among those identified as significant, compared to the general read mapping. Such a difference is most likely caused by introgressions and/or presence/absence polymorphisms that are frequently the origin of resistance to powdery mildew [37]. If such an introgressed region is not present in CS, reads containing significantly associated *k*-mers would not map. To investigate the possible origin of the non-mapped *k*-mers, we used 10 additional *Triticum aestivum* genomes [22] as reference genomes. Using this set of genomes, we found that 53% (8,969) of the significant *k*-mers were mapped in all genomes (**Figure 3E**). We considered a *k*-mer mapped when at least one of the reads containing such *k*-mers mapped to one or more genomes. A further 30% (5,003) were mapped in some but not all of the genomes, and the remaining 17% (2,923) did not map to any of the 10 *Triticum aestivum* genomes **(Figure 3E)**. Thus, by adding the 10 genomes as references, we could map 83% of the *k*-mers, compared to 68% when using only CS as a reference. Some of the *k*-mers that do not map to CS map to new regions of interest in other reference genomes. Besides, some *k*-mers also map to regions that have already been detected using CS, but reduce the width and increase the precision of the peak. Taking *Pm4* as an example, we found that when using SYMattis as a reference genome, the associated region perfectly overlaps with the position of the annotated *Pm4* gene, which is not the case for CS (**Figure 3D).** The *Pm4* gene is known to originate from an introgression from tetraploid wheat [26], and blast analysis shows that the gene is present in SYMattis and absent in CS. This explains the higher precision when using SYMattis as a reference genome. Using the *k*-mer approach, we pinpointed narrower peaks, allowing us to define more precisely genomic regions harboring genetic loci of interest (**Figure 3D**).

### Progenitor genomes allow mapping an additional 8% of the significant *k*-mers

Many introgressions derived from wild relatives in bread wheat have been described to contain resistance genes [38,39]. Therefore, to investigate the possible origin of the 17% non-mapped reads, we selected a set of genomes representing close relatives of bread wheat as well as ancestors/progenitors of hexaploid wheat, including *Aegilops tauschii, Triticum turgidum, Triticum urartu,* and *Triticum spelta* (**Figure 1**). By implementing this step, we mapped an additional 60% (1,742) of the 2,923 non-mapped *k*-mers **(Figure 3E,F)**. Combining all reference genomes, we could unambiguously assign a physical position to approximately 93% of the *k*-mers, which is 25% more of the significant *k*-mers compared to using only CS as a reference genome. In summary, our analysis demonstrates the advantage of using multiple reference genomes: starting from the same set of raw genotypic data, an additional 25% of all significant *k*-mers were mapped, resulting in novel associations and improved resolution of associated genomic regions.

### *k*-mer GWAS allows to detect multiple known *Pm* genes present in the Swiss wheat collection

To identify resistance genes acting against multiple *Bgt* isolates, we analyzed significant *k*-mers shared among multiple isolates (**Figure 3B)**. Due to the complexity of displaying association data from multiple isolates across different genomes, we developed a Shiny app, accessible at https://benjiapp.shinyapps.io/Manhattan_plot/. To generate a Manhattan plot, users must follow the following steps: (i) select a reference genome, (ii) choose an isolate, and (iii) specify the chromosome (s) to display. While only one genome can be visualized at a time, multiple isolates can be compared simultaneously by stacking multiple plots. For a more detailed view of an associated region, users can zoom in by specifying the coordinates of a region of interest. An annotated screenshot is shown in **Supplemental Figure 8.**

The largest group of common *k*-mers (n= 5,594) is shared among the isolates ARG_4_2, JPN_CHIKA, GRB_JIW2, CHE_97251, THUN12, and KAZ_1b. These *k*-mers predominantly cluster at the end of chromosome 7A, a region known to harbor the *Pm1* and *Pm60* resistance genes, and is characterized by suppressed recombination [36]. This clustering aligns perfectly with the phenotype of the *Pm1d/a/e* tester lines, which exhibits resistance to the abovementioned isolates and susceptibility to the rest **(Figure 2B)**. Furthermore, *k*-mers from this group also tag other loci of interest, such as on chromosomes 2B and 6B. Extending this to all the reference genomes we found, for instance, that an associated region at the beginning of chromosome 3D was only detected when using Renan, Lancer, and CS as reference genomes. Out of the 34 main regions detected across the 10 reference genomes and the 10 isolates, only 15 are shared in all reference genomes. Such comparative analysis across multiple genomes highlights the limitations of using a single genome reference, as many significant regions would be missed.

The first four groups (groups 1-4) in **Figure 3B** represent the largest groups of common *k*-mers and are formed by a similar set of isolates with minimal variation. Group 5 comprises *k*-mers private to KAZ_1b, while groups 6 and 8 contrast with the first four groups as they contain *k*-mers found significant for isolates TUR_1C, CHE_96224, and IRN_GOR5. THUN12 isolate is the only member common among all these groups. Group 7 includes significant *k*-mers solely from IRN_GOR5 (**Figure 3B**). This pattern correlates well with the correlation matrix between the phenotypes shown in **Figure 2C**.

Interestingly, we observed two distinct, non-overlapping groups of isolates sharing common *k*-mers. One group includes *Bgt* isolates GRB_JIW2, CHE_97251, and KAZ_1b, while the other group comprises *Bgt* isolates TURC_1C, CHE_96224, and IRN_GOR5. Comparing these data with the pattern of *Pm*-tester lines, we found that a combination of multiple *Pm* genes could explain this distribution. In particular, *Pm2a* and *Pm4a* confer resistance to TURC_1C, CHE_96224, and IRN_GOR5, while showing susceptibility to the other isolates, except for JPN_CHIKA. Conversely, various *Pm1* alleles exhibited resistance to GRB_JIW2, CHE_97251, and KAZ_1b. Examining the mapping of *k-*mers from different groups revealed a clear correlation with the expected *Pm* genes. Groups 1-4 predominantly map to the end of chromosome 7, around the *Pm1* region. *k*-mers from groups 6 and 8 almost exclusively tag the *Pm2* and *Pm4* regions. While we detected associated regions for *Pm2 and Pm1/Pm60* in all of the ten reference genomes, the *Pm4* region is not detected in the Lancer genome.

Such grouping together with the mapping to multiple genomes shows that we can detect clear and strong signals from specific *Pm* genes. Zooming into the region of *Pm4* and *Pm2* using SYMattis as reference genome shows that those signals are directly on top of the position of the genes. (**Figures 4A and B**).

**Figure 4:**
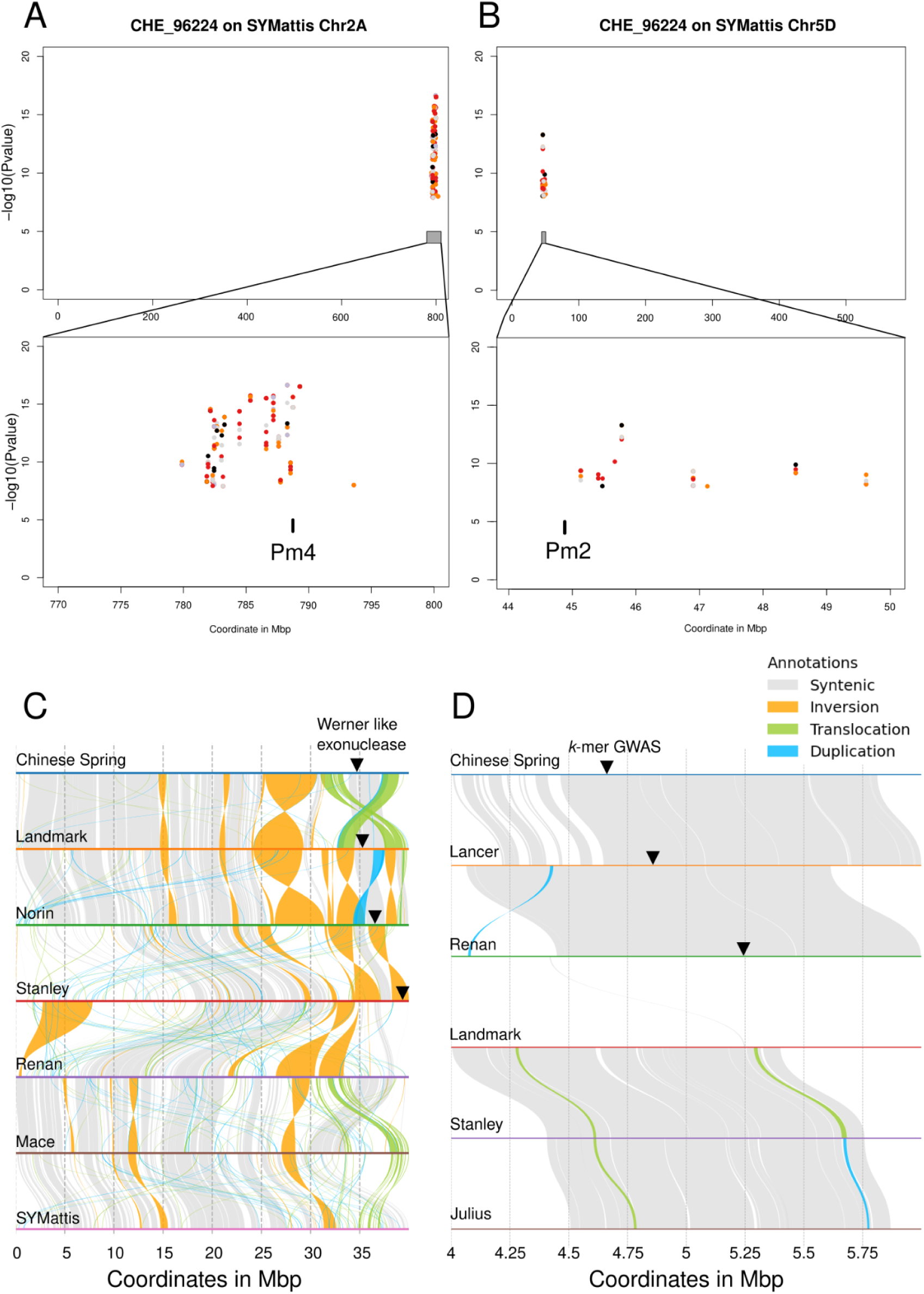
Identification of known *Pm* genes and new candidate genes for powdery mildew resistance. Manhattan plot for the *Bgt* isolate CHE_96224 and the chromosomes 2A (A) and 5D (B) of the SYMattis genome. The zoom-in of each of the regions of interest also shows the position of two of the *Pm* genes known to confer powdery mildew resistance. **C**: Alignment of the genomic regions on chromosome 2B containing the Werner-like exonuclease gene candidate for genomes containing the candidate gene (CS, Landmark, Norin, and Stanley), as well as other genomes not containing it (Renan, Mace, and SYMattis). **D**: Alignment of the genomic region around the candidate gene at the beginning of chromosome 3D (4 to 6 Mb). The position of the GWAS peak, as well as the best gene candidates, are marked with a triangle. A longer alignment of the 12 first Mb of chromosome 3D is in **Supplemental** Figure 18.

### New candidate *Pm* genes identified in the Swiss wheat collection

To discover new candidate genes for mildew resistance, we focused on all regions detected having significant *k*-mer associations not overlapping with previously known *Pm* genes. For those regions, we extracted all protein sequences within 1 Mb around each detected peak. We then used BLASTp for all those genes to infer gene function. After several filtering steps (see Method for details), Manhattan plots for each of the genomes combining the different isolates, as well as a summary table of the main associated regions, are presented in **Supplemental Figure 9.** The table shows the presence of the main regions in the different genomes used as reference. As the genomic coordinates for each genome cannot be directly compared, the correspondence of the region of association between the genomes is arbitrary and some similar regions might not be the same in another genome. For in-depth analysis of candidates of choice, alignment between the genomes of the region of interest will be required.

One intriguing region was identified on chromosomes 3A, 3D, and 2B, associated with resistance against *Bgt* isolates CHE_96224, ARG_4_2, IRN_GOR5, THUN12, and TUR_1C. The associated regions on chromosomes 3A and 3D were only detected when using the reference genomes of Fielder, Julius, and Lancer (**Supplemental Figure 10**). The region on chromosome 2B overlaps with the homologous region associated with *Pm4*, making it challenging to clearly separate the two. The associated regions on chromosome 3A and chromosome 3D directly overlap a Werner-like exonuclease gene (TraesJUL3A03G01434660), present in one and two copies on chromosome 3A and chromosome 3D, respectively. Each of these three copies is perfectly conserved across all the reference genomes used in this study, except for Landmark and Renan, which lack these copies. Due to this conservation across most genomes and the lack of correlation with the resistance patterns of these genomes (**Supplemental Figure 11**), the associated regions on chromosome 3A and chromosome 3D are unlikely candidates for powdery mildew resistance.

Focusing on the association on chromosome 2B, we applied the same approach to examine structural variation across the genomes. To include the entire start of the chromosome we did an alignment of the first 40 Mb of chromosome 2B across all genomes which revealed structural diversity around the Werner-like gene region. In the genomes of CS, Landmark, Norin, and Stanley, this region is conserved, and all these genotypes are susceptible (**Figure 4C**). Importantly, this region is absent in the reference genomes of all resistant cultivars, namely Renan, Mace, and SYMattis. These data suggest that the Werner-like exonuclease gene on chromosome 2B might be a susceptibility gene that has not been previously identified, potentially revealing a new mechanism of powdery mildew resistance. Such a gene has also been found in a (DNA affinity purification sequencing) DAPseq experiment in a search of TaZF binding site. TaZF have been found to be involved in powdery mildew resistance by recruiting both Pm2a and AvrPm2 from the cytosol to the nucleus. [24]

Another region of interest is located at the beginning of the short arm of chromosome 3D, and was only detected when using Lancer, Renan, and CS as reference genomes. Its presence in these three genomes is not consistent with phenotypic observations: while Renan and Lancer are resistant to many of the tested powdery mildew isolates, CS is not resistant to any. To further explore this region, the 2 Mb around the most associated *k*-mer was extracted and aligned with all the genomes. This region is conserved between CS, Lancer, and Renan but not for the other genomes (**Figure 4D**). Given that CS is not resistant, it is expected that haplotype diversity occurs in the region between Lancer/Renan and CS. Indeed, we observed such diversity within the first 500kb of the region, which shows considerable variation between CS and Lancer, but is almost fully conserved between Lancer, Renan and Norin (**Figure 4D**). While Lancer is resistant to 7 of 9 isolates, Renan is only resistant to CHE_96224, IRN_GOR5, THUN12, JPN_CHIKA, TUR_1C, and Norin is susceptible to all isolates. After extracting all the genes annotated in the region from the Lancer genome, we blasted them to all other genomes to find how conserved each of the genes were. The heatmap in **Supplemental Figure 12** summarizes this.

Each of the protein sequences have been also blasted to the NCBI database to extract possible functions. Thirteen genes are found to be NBS-LRR genes, although all of them are fully conserved between Lancer and Norin, including +/- 2kb of the coding region. Because Norin is fully susceptible such genes are likely not responsible for the resistance observed in Lancer. We found 12 genes present only in Lancer (**Supplemental Figure 12**), out of those five are uncharacterized protein, two are Wax Ester synthase like protein, one is a fatty acyl-CoA reductase, one is a glucomannan 4-beta-mannosyltransferase and one is a receptor like prot12. Waxes are part of the cuticle and protect different organs from biotic and well as abiotic stresses [40]. Even though such genes have not been described in resistance to powdery mildew, its presence/absence pattern between the genomes make it a solid candidate. Thus, using *k*-mer GWAS we detected new candidate genes and further analysis of the identified regions and the candidate genes will reveal a possible role of those genes in resistance.

### Using progenitor genomes allows the discovery of new candidates

To find the location of the *k*-mers absent in all tested *Triticum aestivum* genomes we mapped the reads to four progenitor genomes of wheat. The resulting Manhattan plots are represented in **Supplemental Figure 13**. To compare with the location of the known *Pm-*genes we also blasted all the cloned *Pm-*genes to the four genomes and their location is indicated in the Manhattan plot (**Supplemental Figure 13**). In *Aegilops tauschii, Triticum urartu,* and *Triticum spelta*, we detected a region overlapping with the position of *Pm2* or its homolog on chromosome 5A. *Pm2* was originally described as a gene introgressed from *A. tauschii* into chromosome 2D [41]. For all four genomes, we detected a region at the end of chromosome 7 that tags the region of *Pm1* and *Pm60,* originally from *Triticum aestivum*, as well as *Triticum monocucum* and *Triticum urartu*, respectively. For all the other regions we used the approach based on the blast described above to annotate the gene within the 1 Mb around each peak. After filtering the blast results, we ended up with 4, 6, 2, and 15, significantly associated regions containing candidate genes in *Aegilops tauschii*, *Triticum spelta*, *Triticum turgidum*, *and Triticum urartu* respectively. Except for the significantly associated regions on chromosomes 7, 5, and 2, overlapping with *Pm1/Pm60* and *Pm2* and P*m4* respectively, all the other associated regions do not overlap with previously detected regions using the 10 *Triticum aestivum* genotypes as reference genomes. For example, new associated regions were detected at the end of chromosome 5D, 5A in *Aegilops tauschii*, *Triticum urartu*, respectively or at the start of chromosome 3D, 3A, 3A/3B for *Aegilops tauschii*, *Triticum urartu* and *Triticum turgidum* respectively (**Supplemental Figure 13**). Therefore, the use of wild relative genomes as reference revealed additional potential resistance genes undetected in hexaploid wheats, suggesting that resistance genes were left behind or introgressed duringbreeding of hexaploid wheat.

### *k*-mer GWAS allow the discovery of new candidate genes for adult stage resistance to wheat powdery mildew

We evaluated our panel under field conditions for adult plant resistance against powdery mildew over two years. As the panel included spring and winter wheat accessions and the AUDPC (Area Under the Disease Progress Curve) values cannot be compared between these two groups, the collection was splitted in winter (276 accessions) and spring wheat (139 accessions), both showing high R-square coefficients of AUDPC values, 0.68 and 0.87 between the two environments (years), respectively (**Figure 5A**).

**Figure 5:**
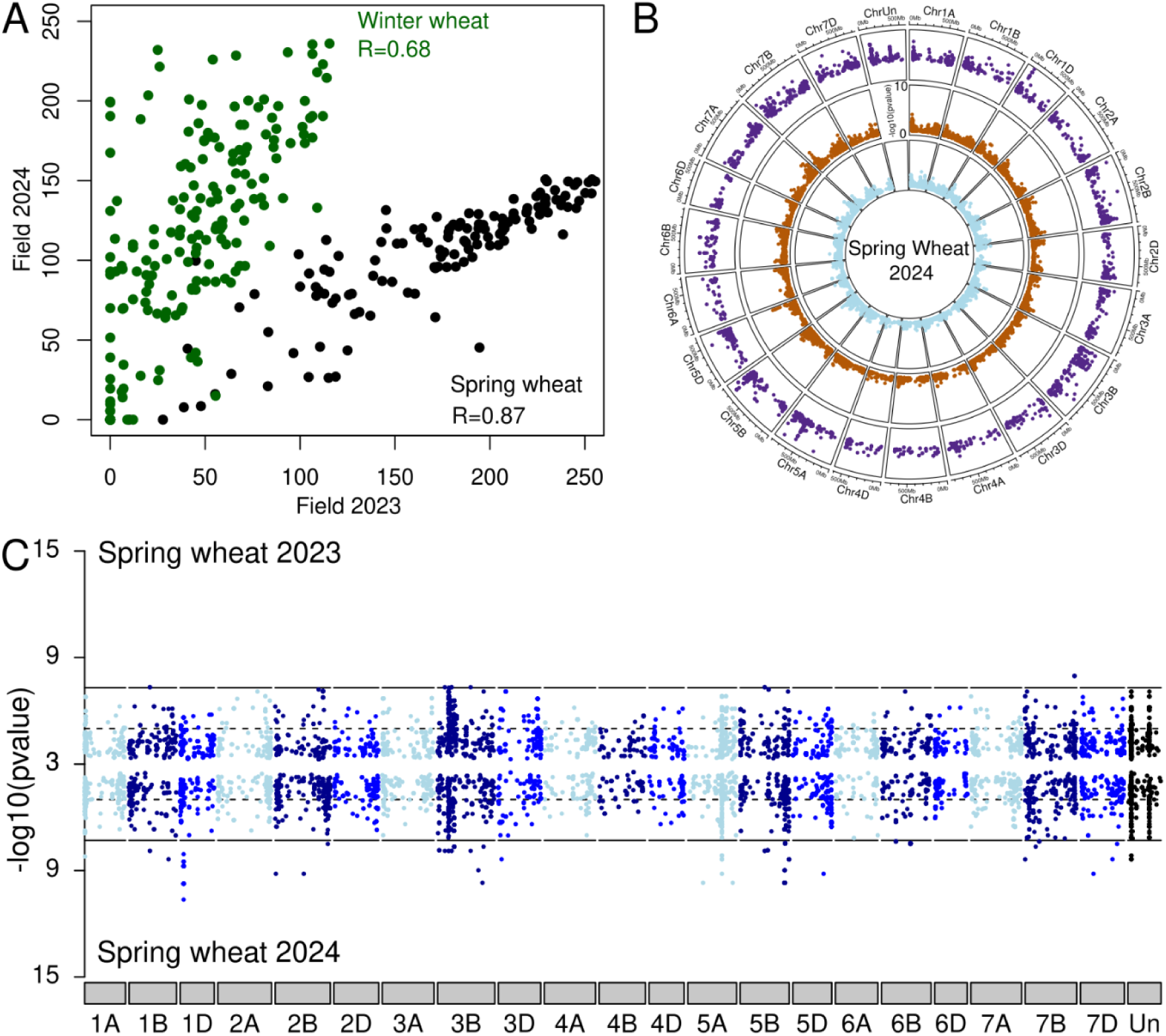
Genetic basis of wheat resistance at adult stage. **A**: Correlation between the two years of field phenotyping for spring (black) and winter (green) wheat. R represents the Pearson coefficient. **B**: GWAS using Spring Wheat 2024 for phenotypic values and for comparison of three genotype matrices. The inner circle is DArTseq SNP, the middle SNPs chip, and the outer *k*-mer GWAS. All SNPs are plotted for the two innermost circles, but only the *k*-mers above -log10(p-value) of 3 are plotted in the outermost circle. **C**: Double Manhattan plot comparing *k*-mer GWAS from the two years of Spring wheat phenotyping. 2023 at the top and 2024 at the bottom. Only *k*-mer with a significance above -log10(p-value) of 3 is displayed. The colors are representing the different subgenomes of wheat. Light, dark, and blue represent the A, B and D genomes.

The identification of genomic regions associated with adult plant resistance in GWAS studies is challenging. Such resistances often involve several genes, reducing their respective association with the observed phenotypes and making it more difficult to be detected by GWAS. To test whether our *k*-mer approach outperforms standard pipelines, we performed a GWAS for adult-stage resistance comparing the *k*-mer and the two SNP matrices using field phenotypes. Here, we focus on the Spring Wheat 2024 trial as an example (**Figure 5B**). We identified a significantly associated region at the end of chromosome 5D (around 500 Mb) using the SNP set obtained through DArTseq and the SNP chip. The associated region on chromosome 5D does not overlap with the region found using the SNP matrices. Moreover, the *k*-mer GWAS identified multiple, narrower associated regions on chromosomes 5A, B, D, and 1D. To compare the GWAS results between two different years we aligned two Manhattan plots together (**Figure 5C**). We found that most regions consistently appeared in both years even though some regions only passed the significance threshold in one of the two years. We observed a similar pattern for the winter wheat data where the same associated regions were detected in both years (**Supplemental Figure 14**). This is expected considering the high correlation between the phenotypes for the two years.

Adding the 10 genomes of *Triticum aestivum* as a reference we again detected additional associated regions present in only one or few genomes. For example, one region on chromosome 4B was only present when Julius’ genome was used as a reference (**Supplemental Figure 15**). We conclude that, despite a limited number of accessions and challenging field phenotyping for adult plant resistance to powdery mildew, using *k*-mers and multiple reference genomes greatly improved the precision and power of GWAS in detecting resistance loci compared to standard pipelines.

## Discussion

To fully leverage germplasm collections for valuable alleles that can enhance crop resilience, comprehensive genetic and phenotypic characterization is crucial—an endeavor that has only recently begun to be undertaken [9,29,42]. However, recovering rare alleles from the vast genetic landscape of global diversity panels can be challenging [43,44]. Alternatively, the use of local or regional diversity panels, better adapted to local agroecosystems [45], may be more suitable for specific breeding applications. In our work here, we address this gap by assembling a collection of 461 wheat landraces and old cultivars, representative of the wheat gene pool in Switzerland, to characterize landscape of resistance to wheat powdery mildew, a persistent disease in Swiss wheat fields and that has evolved across decades [46].

While some association mapping studies in search of all-stage powdery mildew resistance are limited by the use of a single mildew isolate [47–49] or just a few with similar virulence patterns, (Li et al., 2019), our study significantly broadens the analysis by testing the disease resistance to ten wheat powdery mildews from diverse geographic regions, each exhibiting contrasting virulence patterns based on the disease responses of 37 *Pm* tester lines. On average, 80 accessions (17 % of the collection), were resistant to single *Bgt*. This contrasts to similar studies evaluating seedling resistance to wheat powdery mildew. For example, Li and colleagues [50] tested 1292 accessions, where only 4% were resistant to the tested *Bgt* isolate, or [48], who reported overall susceptibility at the seedling stage among 8,316 winter wheats of the German Federal ex situ gene bank. Our results underscore the suitability of our collection as a source of resistance to wheat powdery mildew.

By comparing response patterns derived from the interaction between wheat landraces and the set of 10 *Bgt* isolates with known virulence/avirulence spectra based on 37 *Pm* tester lines, the presence of *Pm1*, *Pm2*, *Pm4,* and *Pm60* resistance genes could be postulated. Further, the presence of *Pm2* and *Pm4* could subsequently be confirmed using diagnostic molecular markers. Of note, the 28 landraces fully resistant to the 10 tested *Bgt* isolates have the same pattern as the *Pm13* and *Pm36* tester lines. Both *Pm2* and *Pm4* genes have been recently cloned and encode kinase-fused resistance proteins [51,52], a new resistance gene family private to *Triticeae* that provides resistance to different fungal diseases in major cereal crops [53]. Further work is needed to confirm if *Pm13* and *Pm36* resistance genes are responsible for the “all isolate” resistance observed in the 28 landraces, or if novel, unknown *Pm* genes are causing this broad-spectrum resistance. If the presence of these resistance genes is confirmed in hexaploid wheat landraces, they could be used more broadly in breeding programs. *Pm13* was cloned from *Ae. longissima* [51], while *Pm36* presence has only been documented in a few wild emmer wheat accessions [52], making its practical use in breeding programs challenging. As landraces are sexually fully crossable with bread wheat, these resistance loci could be directly cross-bred into modern cultivars, sidestepping long and laborious backcrossing with wild relatives like *Ae. longissima*.

As the bimodal-like distribution of seedling response to wheat powdery mildew suggested the presence of major genes controlling resistance, we performed a GWAS with genotyping data generated with the SNP-chip produced by TraitGenetics. Association mapping analysis revealed 53 significant SNPs, pinpointing seven genomic regions associated with all-stage resistance, two of them corresponding to the *Pm1* (or *Pm60*) and *Pm2* resistance genes. This apparent limited capacity of detecting resistance loci may be explained by the intrinsic nature of the SNP-chip. First, the chip can only detect SNPs. However, SVs such as insertions, deletions, duplications, CNVs, and translocations have been reported to underlie disease resistance phenotypes [54]. Second, SNP-chip design relied on a single reference genome, Chinese Spring, and a few accessions, limiting the SNP set that can be detected [55].

To overcome these limitations, we further genotyped the collection with DArTseq technology, a genotyping technology that is not biased by reference genomes, as it sequences restriction fragments produced by restriction enzyme-mediated genome complexity reduction, which theoretically increases its capacity to detect loci of interest compared to the SNP-chip [13]. Using such data and CS as a reference genome, a SNPs matrix containing 10,068 SNPs was generated. Additionally, a matrix of 11,252 polymorphic SNPs was generated using 12k Illumina Infinium 15K wheat SNP array (TraitGenetics GmbH, Gatersleben, Germany). The use of these matrices resulted in significant SNPs associated with wheat mildew resistance overlapping with the presence of known *Pm* genes like *Pm2* and *Pm4* (Figure 3) [26,27]. Interestingly, although the SNPs matrix derived from DArTseq contained less SNPs compared with the SNPs chip, it resulted in sharper peaks. Such associated regions contain SNPs with a high proportion of missing data and only when including SNPs with 80 percent missing data, all SNPs were included for GWAS. This is not surprising knowing that wheat genomes contain many introgressions [39] that would result in such a pattern.

In order to avoid the bias of the reference genome, we generated a matrix of *k*-mers from the raw data of DArTseq. This led to 176 million unique *k*-mers of length 31bp. Such *k*-mers can tag SNPs, as well as other structural variations [21], and it is known that, especially in plant genomes, an important part of genetic diversity is due to structural variations in different forms, such as presence/absence variants (PAVs), CNVs, insertions or deletions [56]. The disadvantage of using *k*-mer is that the location in the genome is not known. When using Illumina sequencing data, reads containing *k*-mers can be assembled into larger segments and subsequently Blast or sequence alignment can be used to find the corresponding gene [21]. However, the nature of DArTseq data does not allow such an approach. To circumvent this limitation and being capable of comparing with the SNPs GWAS, we mapped the DArTseq raw reads to CS. We could detect all the associated regions already detected using SNPs matrices as well as new ones on chromosomes 2A and 5D. However, only 68% of the *k*-mers could be mapped using the reference genome Chinese Spring. The discrepancy with the number of reads mapping (97.3%) again points towards the fact that introgressions are well-known to harbor genuinely new alleles of agronomic interest [37]. To overcome this, we used multiple reference genomes; ten bread wheat genomes, and four genomes from the progenitors of wheat to capture as much variation as possible. Still, the main disadvantage of *k*-mer-based GWAS approaches is the lack of standardized methods for conducting such analyses. This is in sharp contrast to user-friendly programs like TASSEL [57] or GAPIT [58] that have been developed for SNPs-based GWAS pipelines. To apply the *k*-mer-based association mapping analysis to multiple genomes we developed a pipeline. With this, we increased from 11,570 significant *k*-mers when using CS as a reference genome to 15,712 significant *k*-mers. This translated into the identification of 27 additional new regions associated with resistance to powdery mildew. The inclusion of more reference genomes allowed us to map 25% more *k*-mers reaching 93% of all the significant *k*-mers detected using GWAS (**Figure 3**). Among the 27 regions, we focused on two of the most promising candidates: a gene annotated as a Werner-like exonuclease and a Wax Ester synthase gene in chromosomes 2B and 3D, respectively. Notably, none of the regions were revealed when using the SNP matrices, which highlights the higher discrimination power of the *k*-mer-based approach. This also suggests that these gene’s presence or absence may be attributed to introgressions.

In humans, Werner-like exonucleases have been shown to degrade DNA in a structure-dependent manner [59]. In *Arabidopsis thaliana* Werner-like exonucleases have been shown to interact with the Ku heterodimer which is required for the non-homologous end joining pathway of DNA repair [60]. However, no studies to date report their involvement in plant immunity. The correlation with the phenotype shows that the presence of the gene is associated with susceptibility to four *B. graminis* isolates, CHE_96224, IRN_GOR5, JPN_CHIKA, and TUR_1C as well as THUN12 (Figure 5D). The genomes of CS, Landmark, Norin, and Stanley contain the two copies of the Werner-like exonuclease gene on chromosome 2B, while the other genomes do not have any. Susceptibility factors are rare, but some prominent examples have been described in cereal immunity. For instance, the well-known natural mutation of the *Mlo* gene in barley provides broad-spectrum resistance against barley powdery mildew [61], while the *pi21* recessive mutation is linked to long-lasting resistance to rice blast [62]. More recently, genome-edition of the wheat *Mlo* orthologue has proven to confer resistance to wheat powdery mildew without growth penalties [63], while the inactivation of a target for rust effectors, the wheat kinase *TaPsIPK1*, confers broad-spectrum resistance to rust fungi. *CRISPR*/Cas9 mutations of the two copies of the gene on chromosome 2B would clarify if the Werner-like gene is a susceptibility factor, which if proven, would be the second report of a disease susceptibility gene to wheat powdery mildew, opening up new avenues to control the disease.

The region around 5Mb of chromosome 3D is present in four out of all the genomes tested, namely CS, Renan, Norin and Lancer. Such patterns correspond to a possible introgression that is strongly associated with resistance to powdery mildew. As the resistance pattern observed on Renan genome can be fully explained by the presence of a functional allele of *Pm4,* and CS as well as Norin are fully susceptible, we focussed on genes solely present in Lancer. We found that 3 of the 12 genes present in Lancer in these regions are involved in Waxes synthesis. Evenso they are not known to be directly involved in resistance to powdery mildew, waxes of the cuticle are forming a physical and chemical barrier to biotic and abiotic stresses [40]. Further functional and molecular analyses are required to confirm the possible role of such genes in plant defense.

Adult plant resistance to diseases is genetically complex, usually controlled by several genes associated with genomic regions called quantitative trait loci (QTL), resulting from the effect of each QTL acting —with small or larger effects— leading to a partial phenotypic resistance. Detection of minor-effect QTLs is highly challenging [7]. To further test the detection power of our *k*-mer-based GWAS approach over the SNP-based approach, we performed association analysis using adult plant resistance to wheat powdery mildew collected under field conditions over two seasons. We found that when using SNP matrices, no significantly associated regions to mildew resistance were detected, while the use of our *k*-mer GWAS allowed the detection of multiple regions of interest. With our approach, we show that the capacity to detect those small QTLs is achieved compared to the commonly used SNP-based approaches. Implementing our pipeline to future field studies would result in more accurate and larger detection of QTLs, allowing breeders to accumulate major- and minor-effect loci that can be used for the improvement of wheat powdery mildew (and virtually any other disease) resistance.

The *k*-mer-based pipeline presented here offers several advantages over conventional SNP-based association mapping analysis. First, *k*-mers allow for the assessment of almost all types of variations [21]. While SNP matrices can only detect SNPs, *k*-mers can interrogate diversity given by SVs, such as large indels or introgressions. This led to the discovery of loci associated with resistance that would have been missed using SNP matrices. Second, by utilizing *k*-mers directly from raw sequencing data, we could bypass error-prone stages of variant discovery and genotyping, thereby facilitating the identification of causal variants. Third, the inclusion of multiple reference genomes notably expanded the number of potential loci associated with resistance by interrogating broader diversity [20]. All these improvements translate into greater efficiency on the same phenotypic and genotypic basis, both expensive and labor-intensive to get.

With the increase in the number of genomes being sequenced, we foresee that our approach can become very popular as pangenome GWAS using SNPs has not yet been implemented. Our k-mer GWAS approach proves to be the best choice as such markers can more accurately represent genome diversity, and detailed comparison between available reference genomes would lead to the identification of useful alleles and allele stacking strategies to sustain resistance breeding activities.

## Methods

### DNA extraction and genotyping

The 461 accessions from the Swiss collection have been genotyped using Diversity Arrays Technology Pty Ltd (http://www.diversityarrays.com/) for sequencing and marker identification as a batch of the AGENT project. Seeds were provided by the Swiss gene bank at Agroscope. Individual plants for DNA extraction were grown in a climate chamber cycled at 20 °C/16 °C, 16/8 h photoperiod with 60% relative humidity. Two segments of the first leaf (each approximately 3 cm long and 0.3 to 0.5 cm wide) were placed in 2.2 ml tubes with two 4mm glass beads (ROTH). Samples were frozen in liquid nitrogen and ground with a Geno/Grinder (SPEX SamplePrep) at 1500 rpm for 1 minute. Subsequent steps for DNA binding, washing, and elution were carried out using the automated KingFisher™ Apex Purification System (ThermoFisher) as described in [64,65]. Purified DNA was dissolved in 10mM Tris HCl pH 8.0 and shipped frozen with blue ice.

### SNP chip generation

Plants were genotyped using an Illumina Infinium 15K wheat SNP array (TraitGenetics GmbH, Gatersleben, Germany) composed of 13’006 SNPs. The sequences and the position of the molecular markers on the IWGSC CS RefSeq v2.1 were retrieved from the 90K iSelect (Kansas State University, Manhattan,USA) and the Breeders’ 35K Axiom® arrays (Axiom, Santa Clara, USA) from which originated the Illumina Infinium 15 wheat SNP array. This led to unambiguous positioning of 11,983 markers out of the 13’006 SNPs. 1,023 markers remained with unclarified physical position, either because (i) originally not mapped to any chromosome (41 markers), or (ii) mapped to two or more positions often on the three wheat homoeologous chromosomes (982 markers). These unmapped or not precisely mapped markers were blasted against the IWGSC Cs RefSeq v2.1 using GrainGenes online tools (available at: https://wheat.pw.usda.gov/GG3/, accessed 12/08/2024). The physical location associated with the lowest E-value was retained. This leads to unambiguous positioning of 981 additional SNP on the reference genome sequence representing a total of 12’964 SNPs Moreover, 415 SNP markers containing more than 25% of missing information were trimmed-off the SNP matrix. The remaining SNPs table was imputed using Beagle 5.4 [66] and 733 SNPs with a minor allele frequency (MAF) of less than 5% were removed. After the different cleaning operations, 882 SNPs (on a total of 11’816 SNPs) could be reassigned by BLASTING to unambiguous position IWGSC CS RefSeq v2.1 and could serve as a genotypic table obtained after 15K Chip genotyping.

### Mapping

The fastq files containing the sequencing data from DArTseq were mapped to the different genomes using BWA mem [67]. While duplicated reads were marked by Markduplicates from Picard (v1.101; http://broadinstitute.github.io/picard/), samtools (v1.13) was used for file format conversions like sorting and indexing [68]. The transformation from the obtained “.bam” files to “.bed” files was done using BEDtools v2.30.0 [69]. Such files were further processed for different analyses using R. The scripts are available on GitHub.

### Read processing and Variant calling from the DArTseq data

Adapter sequences from raw reads obtained from DArTseq sequencing were trimmed using cutadapt (v1.9.1) [70] with a minimum read length of 30bp. Reads were aligned against the hexaploid wheat reference genome assembly cv. Chinese Spring (RefSeq v2.1) [32] using BWA-MEM (v0.7.15) [71] with default parameters and the output was converted to binary alignment map (BAM) format using SAMtools (v1.3) [68]. BAM sorting was performed using NovoSort (V3.06.05). Variant calling was performed using the mpileup and call functions with the multiallelic-caller (-m) from BCFtools (v1.12) [72] with a minimum read quality (-q) cutoff of 20 and retaining allelic depth and the number of high-quality bases (-a AD,DP) for variant sites. SNPs were further filtered for minimum QUAL ≥ 40, minimum read depth for homozygous calls ≥ 2, minimum read depth for heterozygous calls ≥ 4, and a minimum presence rate of 80% using a custom awk script.

### Powdery mildew isolates genetic analysis

The raw Illumina reads were filtered, mapped, and haplotype-callled to the CHE_96224 reference genome as previously described [23]. Following the same methods from before, the generated SNPs were filtered with vcftools (v0.1.16) with the following parameters:--minDP 3, --maxDP 1000, --maf 0.01 (minor allele frequency), --max-missing 0.999, excluding all SNPs on chromosome Unknown, removing InDels and keeping only biallelic SNPs. In order to look at the genomic diversity of the wheat powdery mildew isolates we performed a PCA on all the available isolates and then excluded the populations for which we did not eventually choose any representatives (i.e. CHN, AUS, and USA wheat mildew populations), resulting in 265 isolates. We performed the PCA analysis of these isolates using vcftools (v0.1.16). We then visualized the PCA via the R packages tidyverse (v2.0.0), ggplot2 (v3.5.1), ggrepel (v0.9.5), and rcartocolor (v2.1.1) [73–76]. The raw sequences of the datasets used in this study can be found in the SRA (Short Read Archive) of the National Center for Biotechnology Information (NCBI) under the accession project numbers found in the papers: [23,77].

### The Swiss collection and *Pm* tester lines

The Swiss collection consists of 461 accessions: 139 are classified as spring wheat and 276 as winter wheat the rest are of unknown growth habit. Accessions were named after villages in Switzerland. Their location of each accession has been extracted based on the corresponding village name in Switzerland. These are approximations, as no precise locations are available for those accessions. Full details of the accessions are available in **Supplemental Table.** The set of tester lines consists of 37 different lines that carry 24 different *Pm* resistance genes.

### Powdery Mildew isolates and phenotyping infections

The following powdery mildew isolates have been used in this study: CHE_96224, KAZ_1b, JPN_CHIKA, GRB_JIW2, ARG_4_2, TUR_1C and POL_3 are described in [23]. Additionally, **CHE_97251** [26], **THUN12** [78], **IRN_GOR5**.

The infection of the leaf segments as well as the scoring was done as described in [29]. The infection score represents the percentage of the leaf segment covered by powdery mildew and varies between 0 and 100. Accessions were considered resistant with a score below 20 and susceptible with a score above 20. Some examples of samples are shown in **Figure 1C**.

### Field phenotyping

Field setup and phenotyping were performed as described in [79]. All accessions have been sown for Winter wheat 2023: 11.10.22, Spring wheat 2023: 2.3.23, Winter wheat 2024: 11.10.23, and Spring wheat 2024: 26.3.24. In short, each accession has been planted in 4 replicates/blocks of 1.5×1m. Those blocks are randomized across the field. Powdery mildew scoring was performed as in [80] over the spring and summer years 2023 and 2024. In 2023, spring and winter wheat were phenotyped at 6 and 4 time points over 36 and 20 days respectively. In 2024, 5 and 6 time points spanning 21 and 42 days for spring and winter wheat respectively (**Supplemental Figure 16**).

### Genomes used as references

For each of the 10 *Triticum aestivum* as well as the 4 progenitor genomes used as reference in this study, the full genome sequence as well as the annotation as gff3, and the protein sequence of the genes have been downloaded from https://plants.ensembl.org/info/data/ftp/index.html. The genomes used are as follows: from *Triticum aestivum*: Julius, Renan, SYMattis, Lancer, Chinese Spring, Fielder, Jagger, Landmark, Mace, Stanley and *Triticum urartu, Triticum turgidum, Aegilops tauschii, Triticum spelta.* A list of all the genomes with the links to the genome sequence/annotation and protein sequence can be found in the **Supplemental Table**.

### Generating *k*-mers

DArTseq produces a high coverage of reads, but for short regions, the first step before generating the *k*-mers was to remove all the perfectly duplicated reads. The program Clumpify (part of the package bbmap) v39.00 [81] was used, with default parameters (zl=9 and dedupe=t). This decreased the number of retained reads by about half. This was done as two identical reads will produce the same *k*-mers.

Based on the pipeline developed by [21], *k*-mers of length 31bp have been generated from the filtered fastq files for each accession using KMC v3 [82]. All parameters have been kept to default except for the -Ci, which sets the threshold for how many times a *k*-mer needs to occur to be counted. Values of 1,2 and 3 were tested (**Supplemental Table**) and similar proportions and associated regions were found for all. We used the 5% family-wide threshold to select the significant kmers. The adapted pipeline can be found on https://github.com/benjj212/Kmer_GWAS_AGENT.

### GWAS

*k*-mer GWAS was performed on 10 traits corresponding to the 10 *Bgt* isolates following the methodology described in [21]. This pipeline uses GEMMA with a Linear Mixed Model, correcting for population structure using a kinship matrix. The default minor allele count (MAC) of 5 was used. For the SNPs GWAS, the GEMMA software was used [83] with option -lmm 2 and -maf 0.05. The Manhattan plots were generated using the package qqman (version 0.1.9) [84].

### *k*-mer analysis

The *k*-mers passing the 5% threshold were used for the 10 isolates, while all *k*-mers were used for the field dataset. These include all *k*-mers that exceed the -log10 threshold for the 5% family-wise error rate [21]. The number of *k*-mers as well as the -log10 threshold for each isolate is presented in **Supplemental Figure 4.** From those *k*-mers, the list of reads names containing the corresponding sequences was extracted from the raw fastq files using the fetch_reads_with_kmers-master from https://github.com/voichek/kmersGWAS. The adapted pipeline is available at https://github.com/benjj212/Kmer_GWAS_AGENT. Using the read names, the bam files were filtered to retain only those specific reads. The filtered bam files were then transformed to BED format using the bamtobed function from the bedtools package [69]. Files containing the sequence of each significant *k*-mer, their p-values, and the coordinates of the corresponding genomes were used for the generation of the Manhattan plot, as well as downstream analyses.

### SyRI

First, the different Fasta sequences were aligned using Minimap2.1 [85] with -ax asm5 -eqx to generate a .sam file. Once the alignments were generated we used SyRI 1.6.3 [86] to generate the comparison files and detect the different types of structural variants between the two sequences of interest. We then used the function plotsr 1.1.1 [87] to generate the output plot. An example of such a pipeline can be found at https://github.com/benjj212/Kmer_GWAS_AGENT.

### Map/plots correlations / Upset plot

All maps and plots were generated using R. Scripts used to generate the plots from the different figures are available on https://github.com/benjj212/Kmer_GWAS_AGENT. For some of the manhattan plot the package CMplot have been used [88]. Shiny app for data visualization:

Manhattan plot: https://benjiapp.shinyapps.io/Manhattan_plot/. Plot for the phenotype distribution: https://benjiapp.shinyapps.io/Map_agent_pheno/. package of the shiny app with all the data are also available on https://github.com/benjj212/Kmer_GWAS_AGENT.

### Candidate gene selection

For each genome, we combined coordinates of reads containing *k*-mers with the P-value from each isolate to detect the main peaks and generate Manhattan plots. Going through the genomes with a window of 1 Mb, all the windows containing at least 10 significant *k*-mers were extracted. Then from each region, all the protein sequences of the annotated genes within the region were saved. Each of the protein sequences were blasted to the plant NCBI database (taxid:3193). The blast has been automated using a Python script available on https://github.com/benjj212/Kmer_GWAS_AGENT. For each blast, the first two results were extracted and listed. The regions known to contain already characterized Pm genes were then tagged. The different steps are graphically explained in **Supplemental Figure 17**.

## Author contribution statement

JSM and BK conceived the project. MH, KG, and BC provided the SNPs matrices. EJ performed the field trials. GH performed the DNA extraction. RL and YL performed the genotyping of the accessions. MB, MPJ, PB performed the library preparation. BJ, YV and MH developed the pipeline. VW and JSM performed infection tests. BJ and AS performed bioinformatics analysis. BJ, JSM and BK analyzed the data and wrote the manuscript. All authors revised the manuscript.

## Acknowledgements

We would like to thank Zoe Bernasconi and Lukas Kunz for their input and ideas in analysis and interpreting the data. We thank the whole field group of Agroscope Reckenholz as well as Elina Leu, Cygni Armbruster, Fabio Gemma, and Lili Hue for their help with the field trials. We thank Wieslaw Podyma and Sylwia Kowalik for their help in the library preparation. The AGENT project has received funding from the European Union’s Horizon 2020 research and innovation programme under grant agreement No 862613. The work was supported by Swiss National Science Foundation grants 310030B_182833 and 310030_204165. B.K. was supported by University of Zurich core funding. JSM is recipient of the grant “Ramon y Cajal” Fellowship RYC2021-032699-I funded by MCIN/AEI/https:// doi. org/ 10. 13039/ 50110 00110 33 and by the “European Union NextGenerationEU/PRTR”.

