## Supplementary Figures for "*k*-mer-based GWAS in a wheat collection reveals novel and diverse sources of powdery mildew resistance"

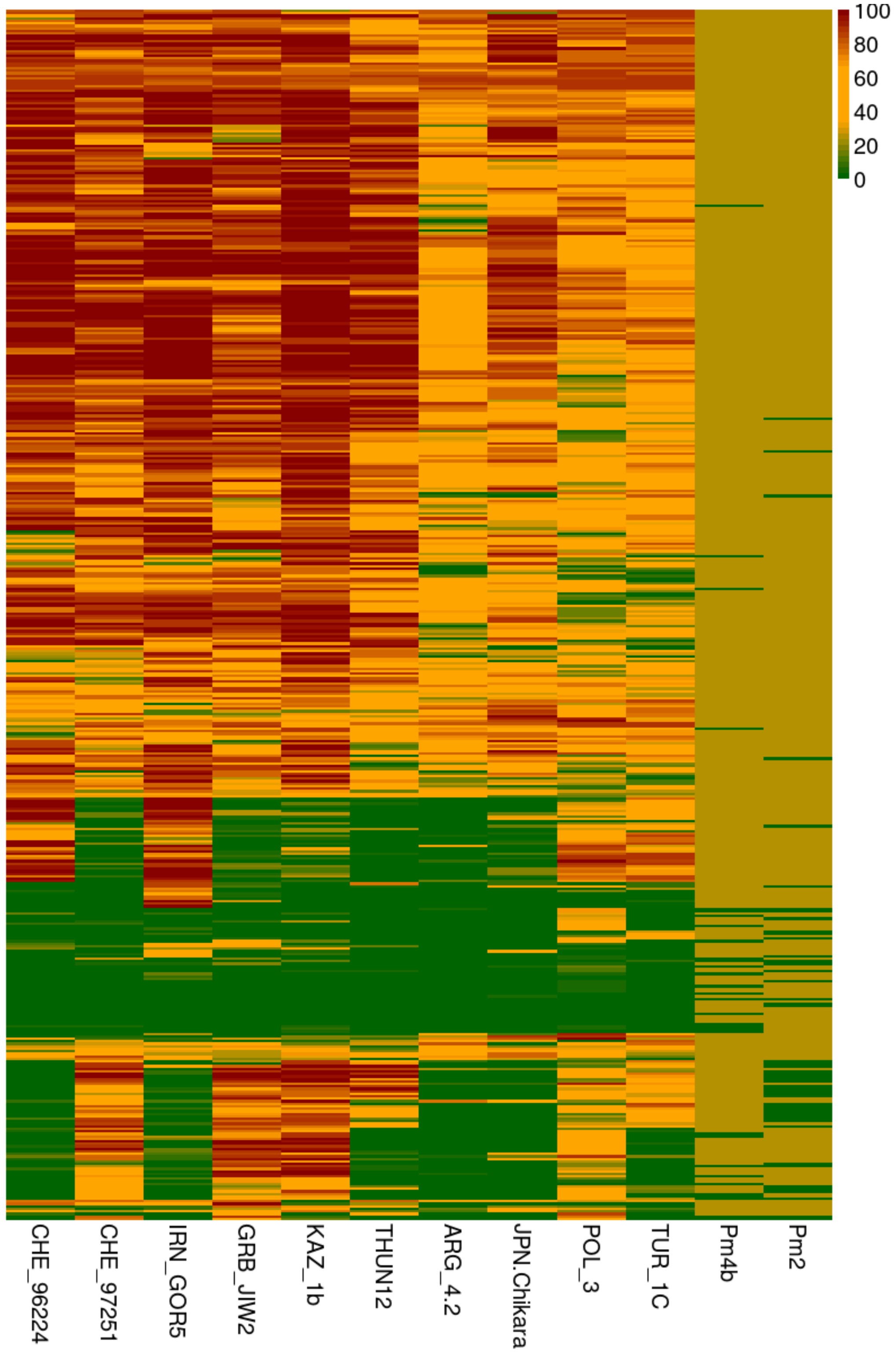


**Supplemental Figure 1:** Heatmap showing the resistance pattern across the 10 *Bgt* isolates for all the accessions of the Swiss collection together with the corresponding genotype of *Pm4b* and *Pm2* for each accession. For the genotyping columns (*Pm4b* and *Pm2)*, green represents the presence of the gene.

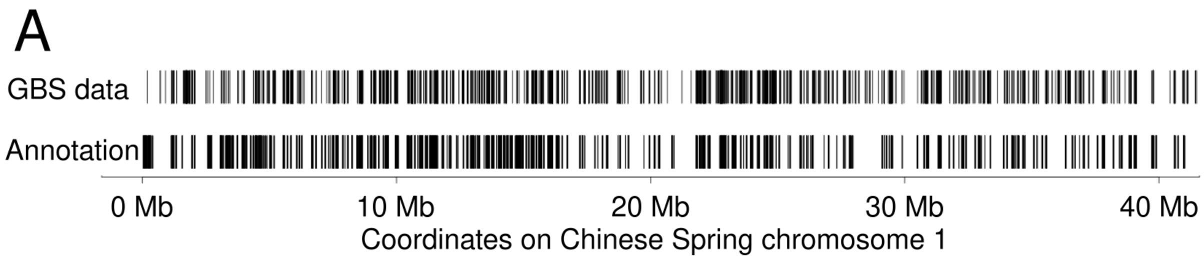


**Supplemental Figure 2:** Evaluation of the mapping distribution of the DArTseq data. A: DArTseq reads mapping to Chinese Spring chromosome 1 compared with the gene annotation of the same genome. B: Proportion of DArTseq mapping within an annotated gene or within +/- 10kb of an annotated gene.

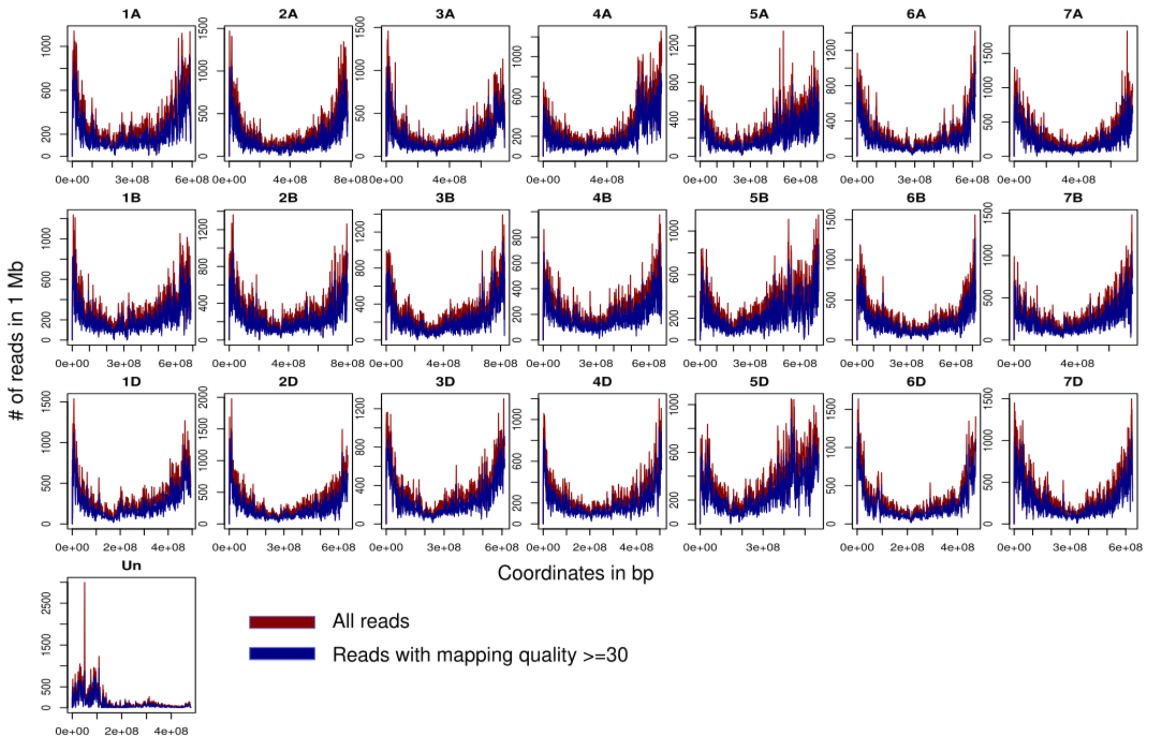


**Supplemental Figure 3**: Profiles of reads mapping density across the 21 chromosomes of the Chinese Spring reference genome. Un: represent the unassembled contigs. All the reads from all accessions were pooled and then mapped to the reference genome.

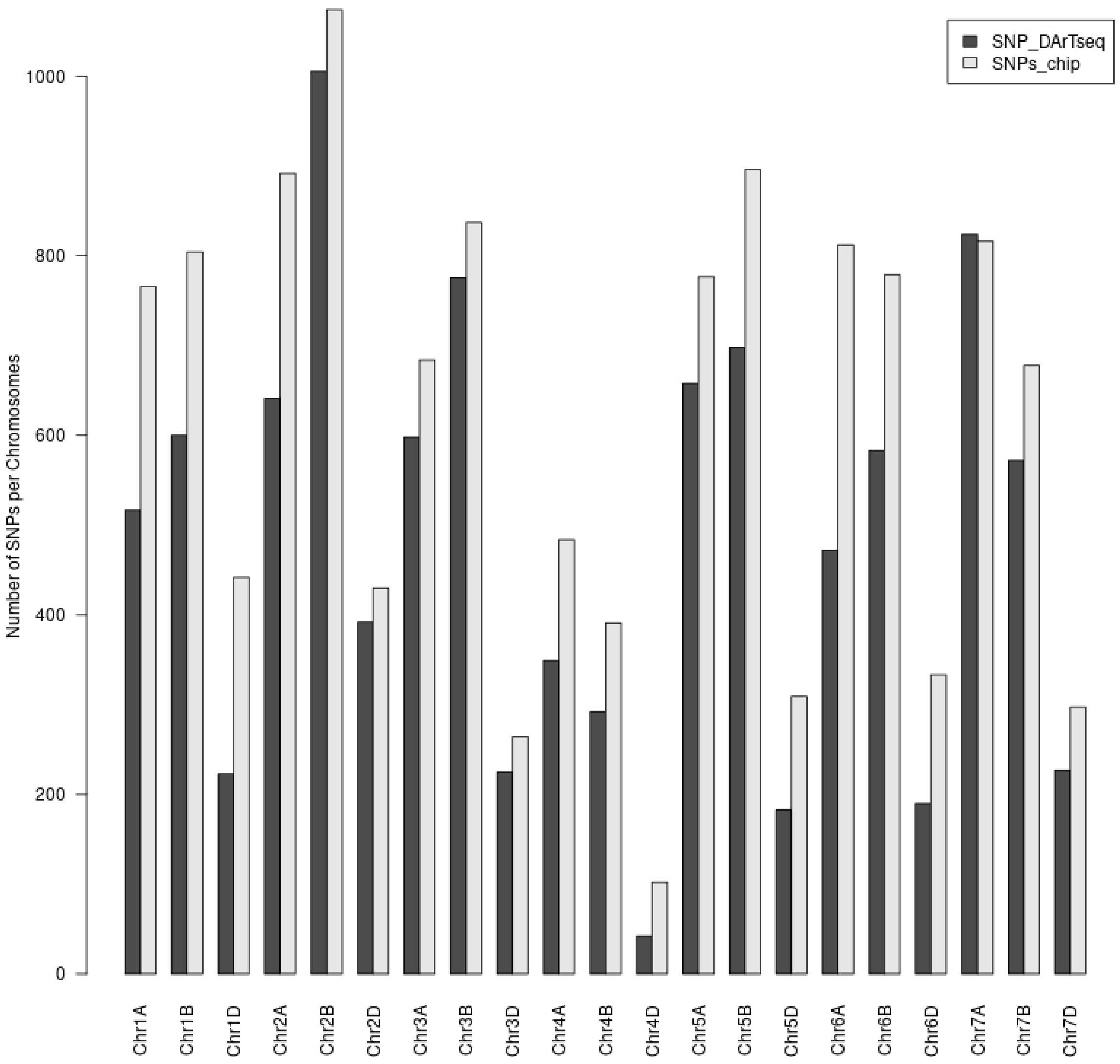


**Supplemental Figure 4**: Barplot showing the number of SNPs per chromosome of the wheat genome for the SNP_chip matrix and the SNP matrix created from the DArTseq data.

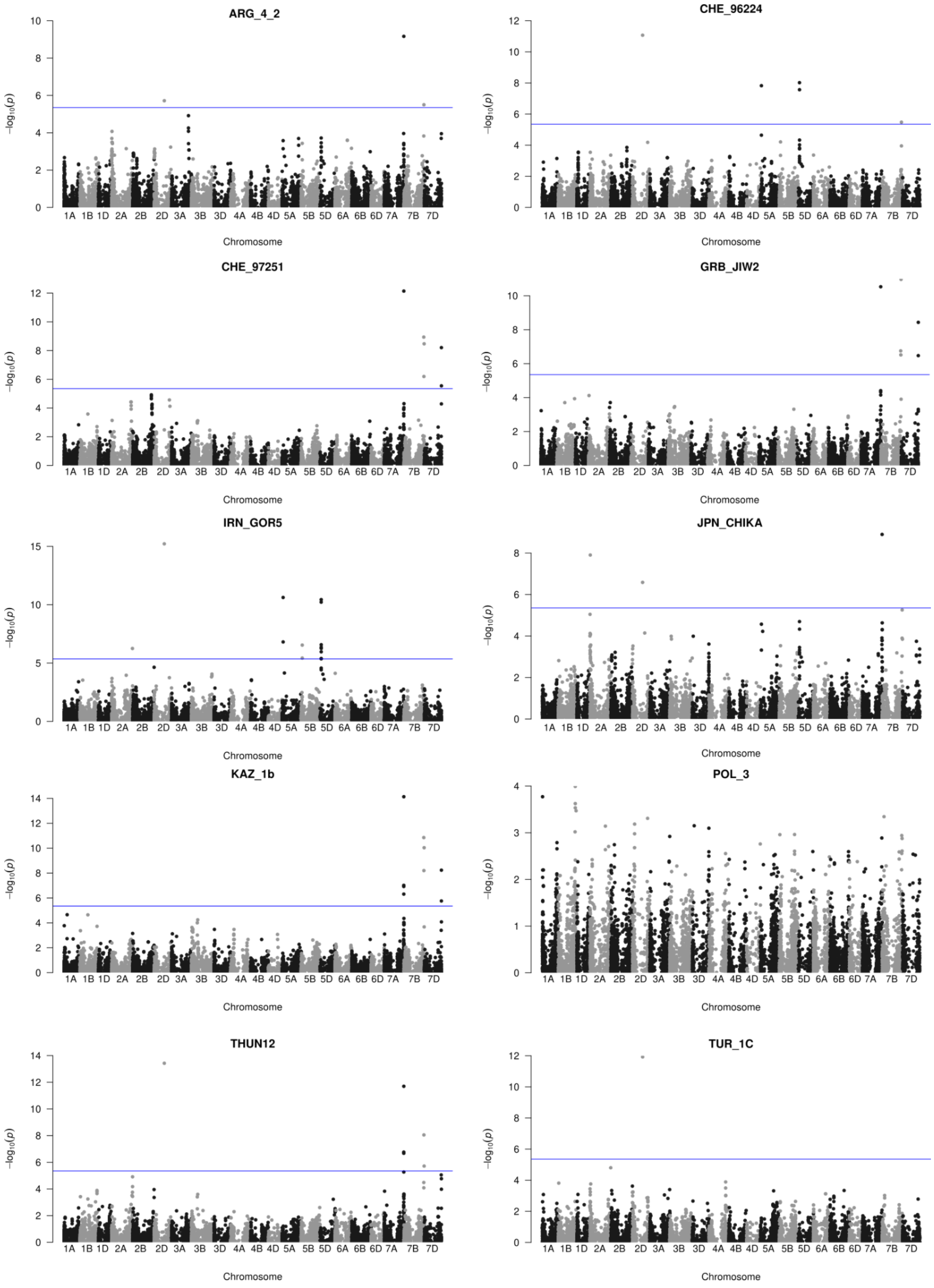


**Supplemental Figure 5:** Manhattan plots for the 10 isolates using the SNP matrix generated using an SNP chip providing the genetic markers for the GWAS. The blue line represents the Bonferroni threshold. The scale is adapted to each plot.

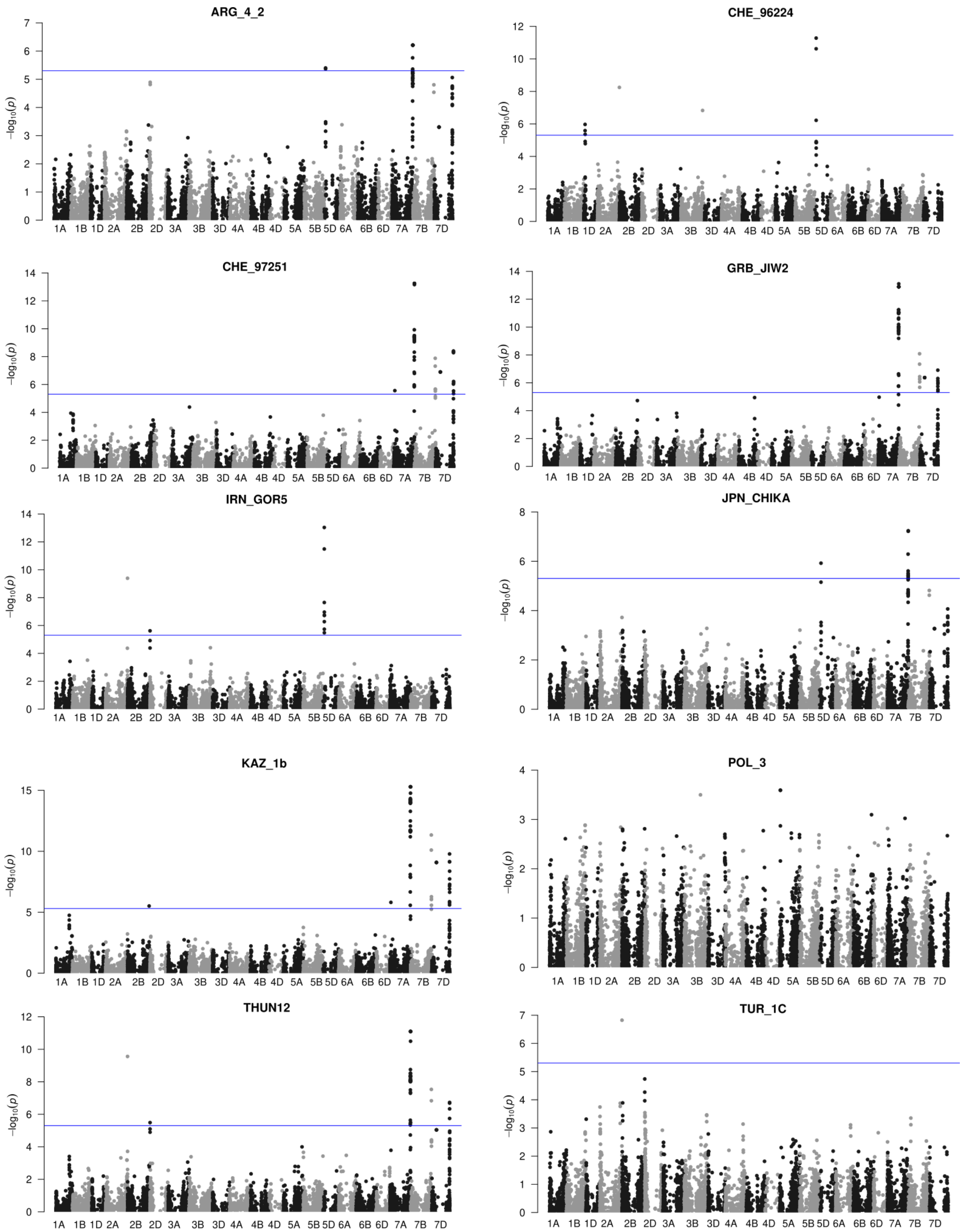


**Supplemental Figure 6:** Manhattan plots for the 10 isolates using the SNP matrix generated from the mapping of the DArTseq data to the Chinese Spring reference genome. The blue line represents the Bonferroni threshold. The y-axis scale is adapted to each plot.

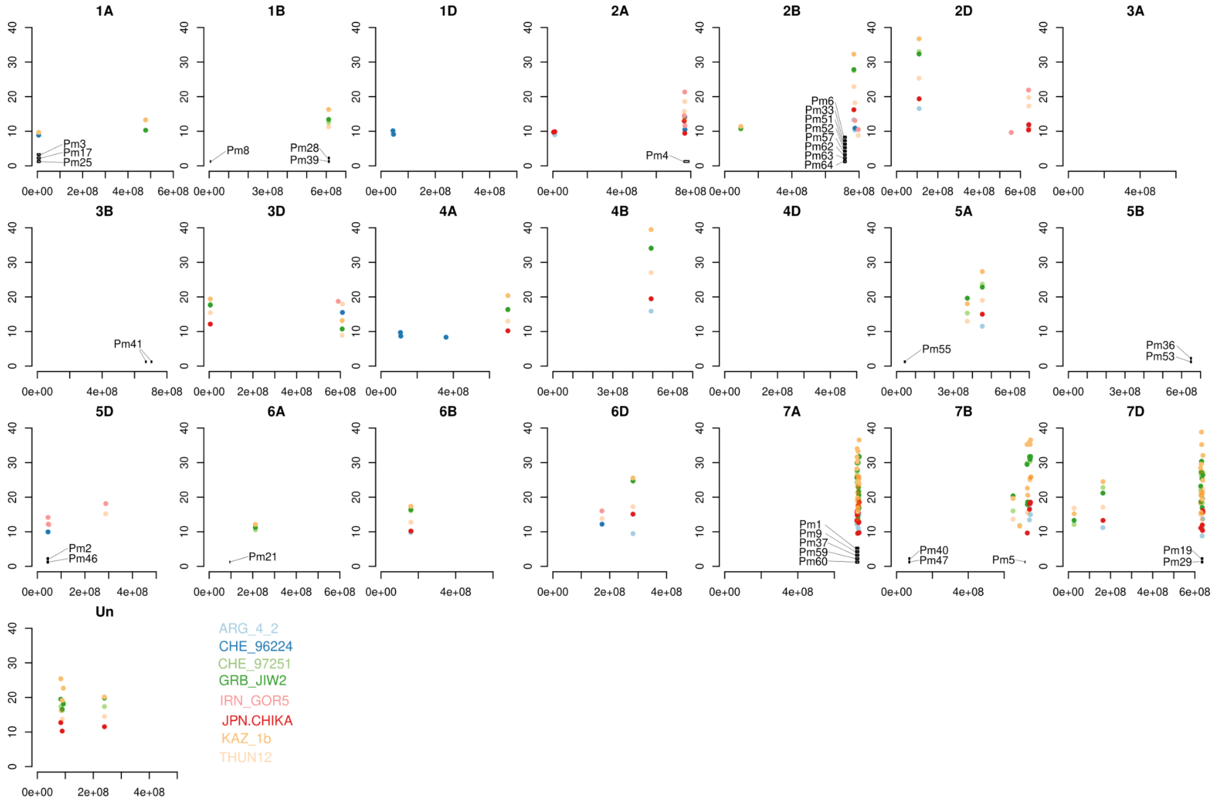


**Supplemental Figure 7:** Manhattan plot from *k*-mer GWAS using the CS genome as a reference. *Pm* gene locations are inferred from blast and previous publications (**Supplemental Table 1**). Colors represent the different isolates.

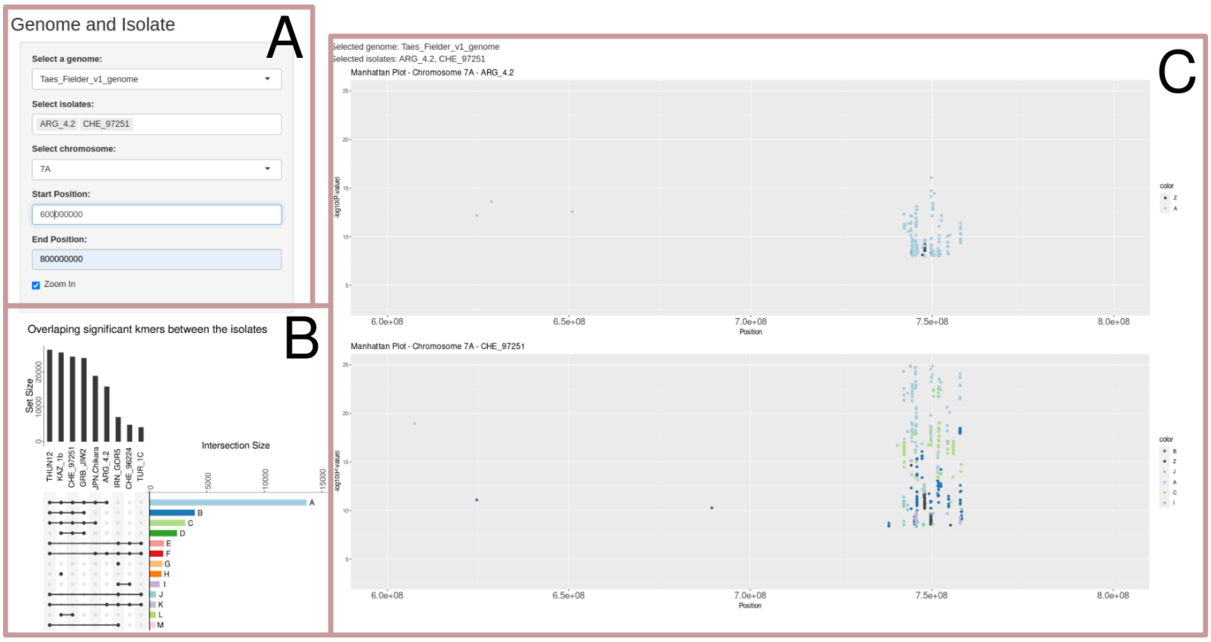


**Supplemental Figure 8**: Screenshot of the Shiny app to explore the GWAS data. Panel A shows the selection of the genome of reference, the isolates to display, and the chromosome. Another option is to zoom in on a specific region by giving the start and the end of the region in bp. Panel B shows the upset plot as in Figure 4A. The different groups are colored as displayed on the GWAS plot (Panel C).

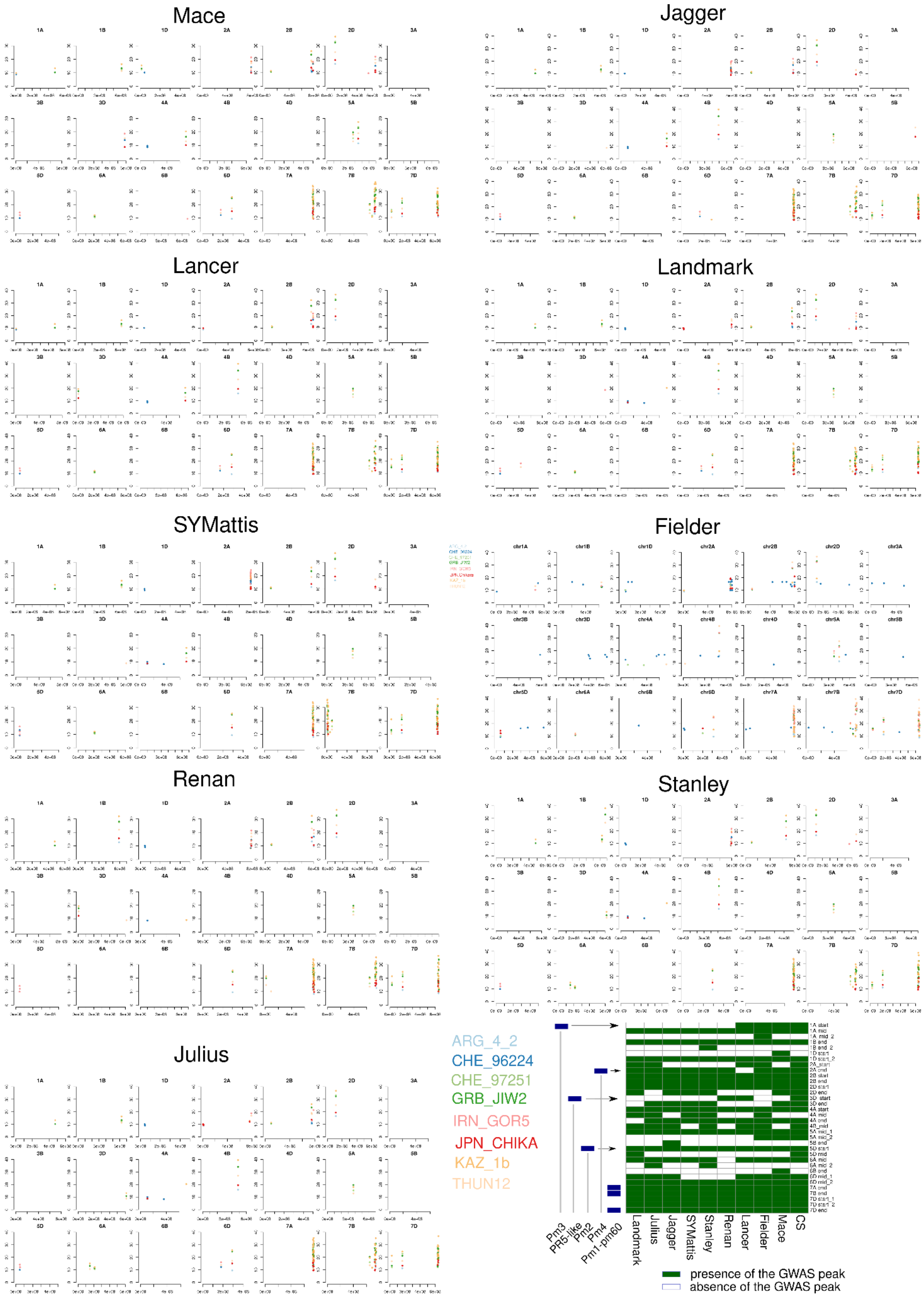


**Supplemental Figure 9:** Manhattan plots of nine reference genomes where different colors represent associations for the various isolates. A summary table describes the presence and absence of the main peaks based on their overall chromosomal positions.

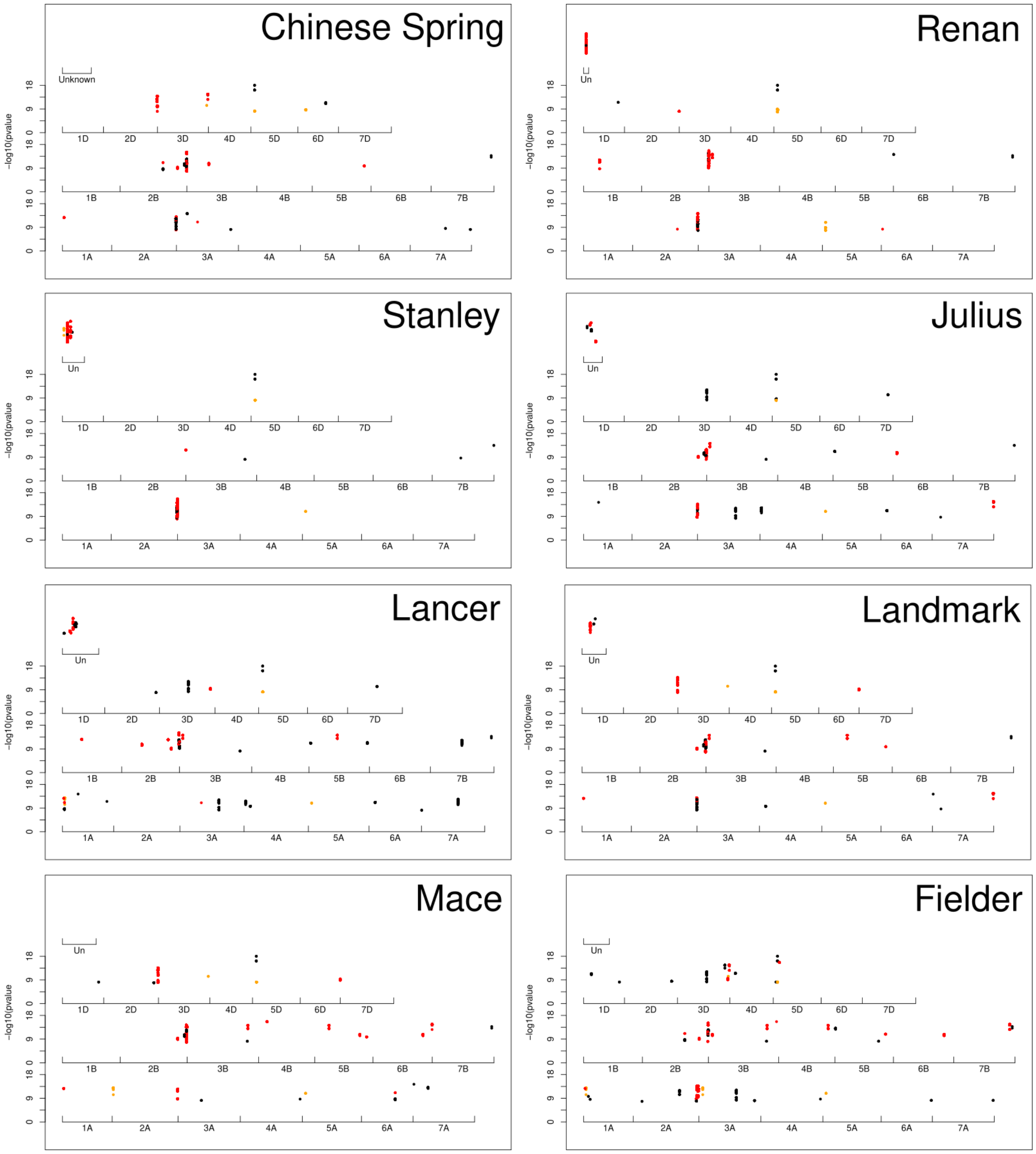


**Supplemental Figure 10**: Manhattan plot from the isolate CHE_96224 using eight different genomes as a reference. Only the significant *k*-mers are plotted. The *k*-mers have been colored based on the result using the SYMattis genome. Red *k*-mer is found to be associated with the region around *Pm4* and orange with the region around *Pm2.* Black was not found in any of those two regions using the SYMattis genome as a reference.

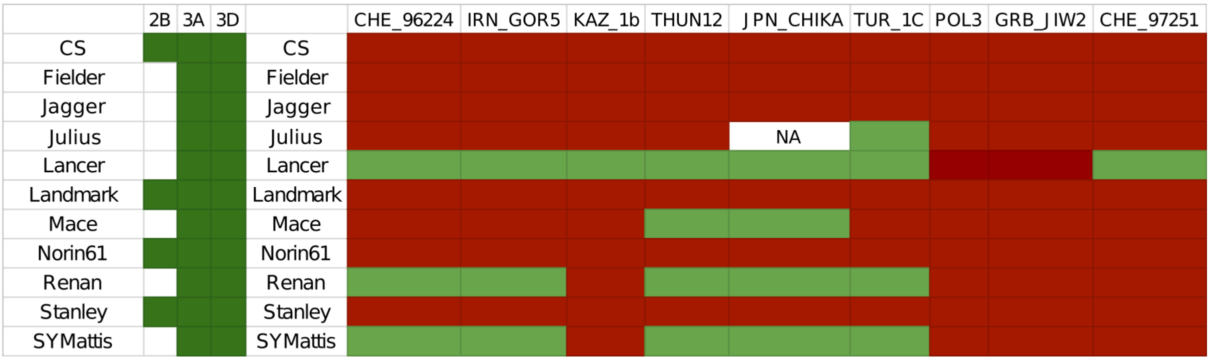


**Supplemental Figure 11:** Table of the presence (green) or absence (white) of the copies of the Werner-like domain on the chromosomes 2B, 3A, and 3D across all reference genomes. The corresponding resistance pattern for each of the genomes across the 10 isolates is represented on the right. Red means susceptible and green means resistant.

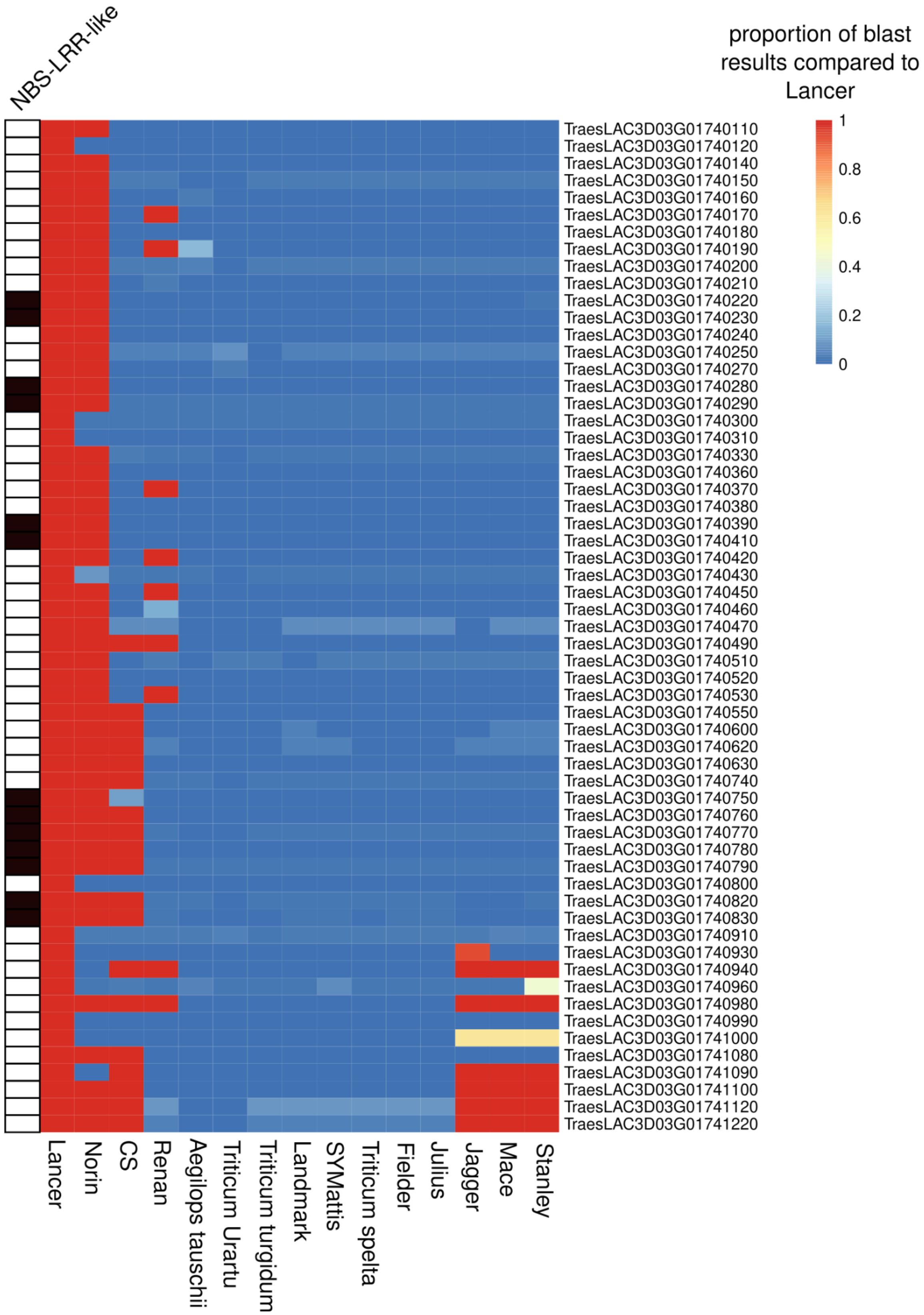


**Supplemental Figure 12:** Heatmap representing the conservation of all the genes within the regions. Genes have been extracted from Lancer genomes and blasted to the other genomes. The color represents the proportion of the gene length that perfectly matches the Lancer gene.

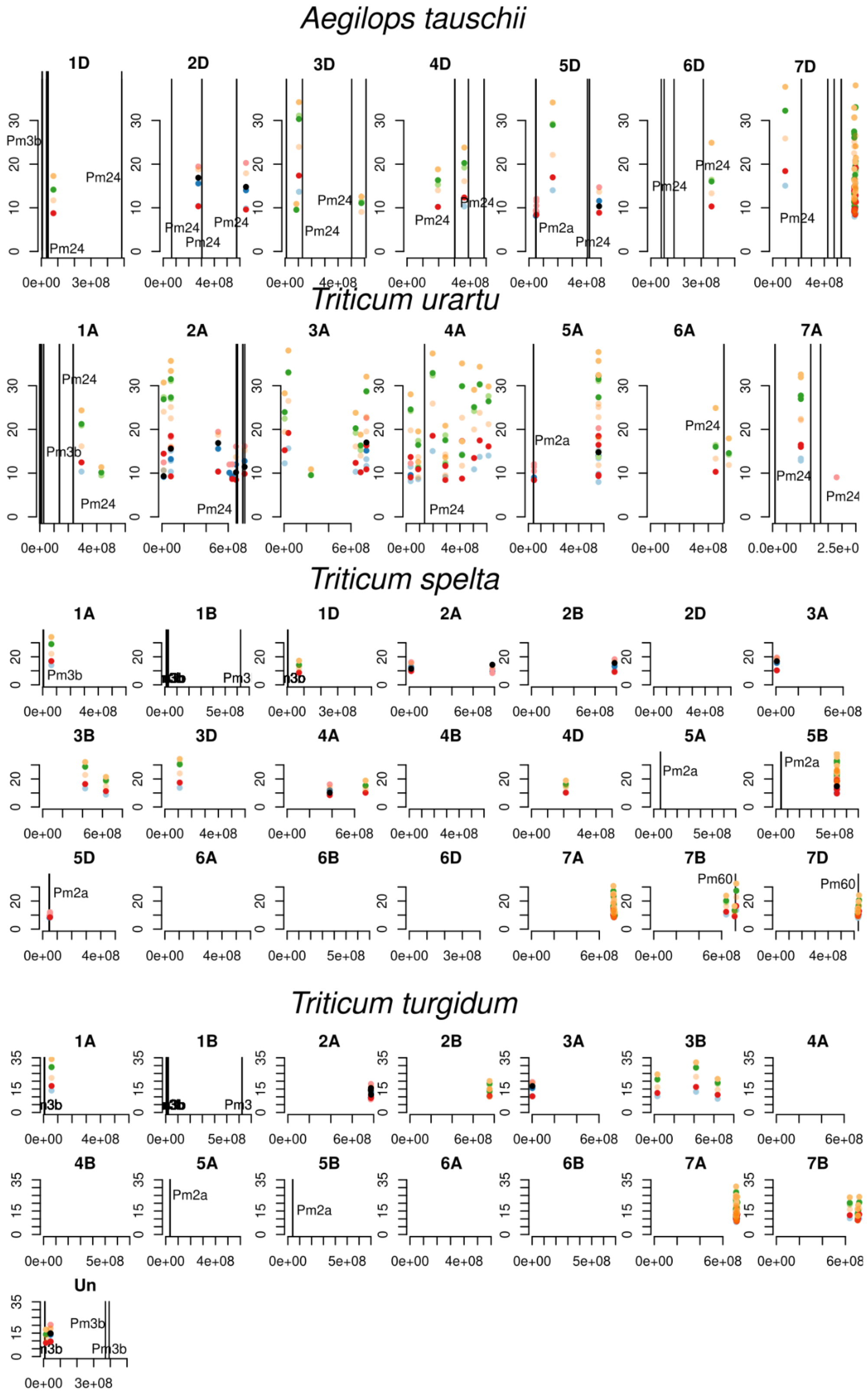


**Supplemental Figure 13:** Manhattan plot using the progenitor genomes as reference. Colors represent from which isolates the significant *k*-mer have been detected. The colors are as in **Supplemental Figure 10**. Additionally, the position of *Pm* genes from blasting gene sequences to the genomes is displayed.

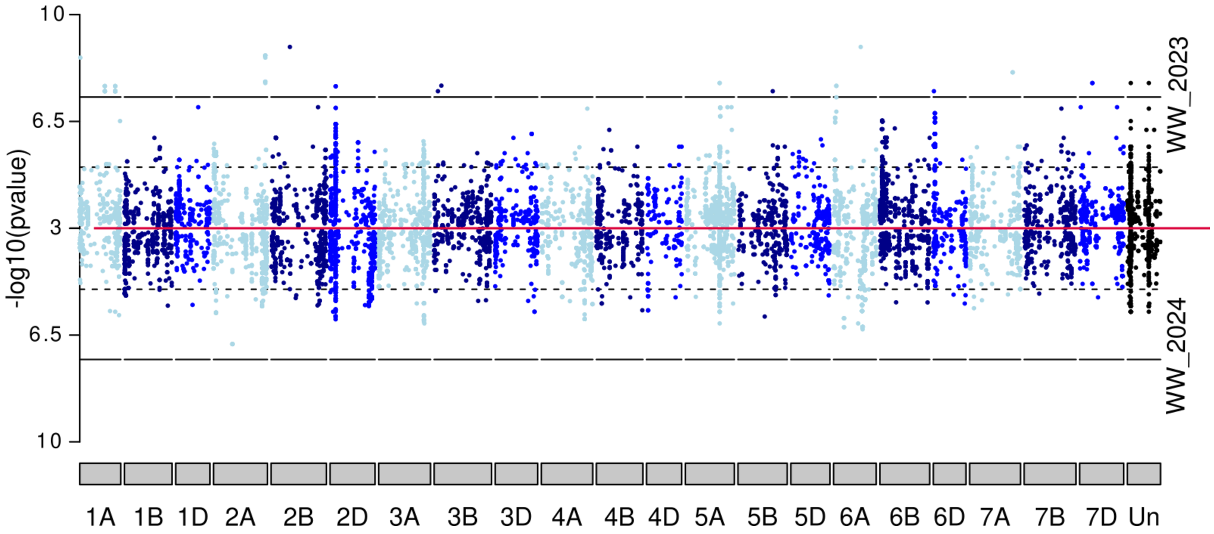


**Supplemental Figure 14**: Manhattan plot comparing field resistance for winter wheat from trials in 2023 and 2024. The -log10(pvalues) have been plotted from 3 onward. The red line separates the two Manhattan plots. This plot is based on the CS reference genome.

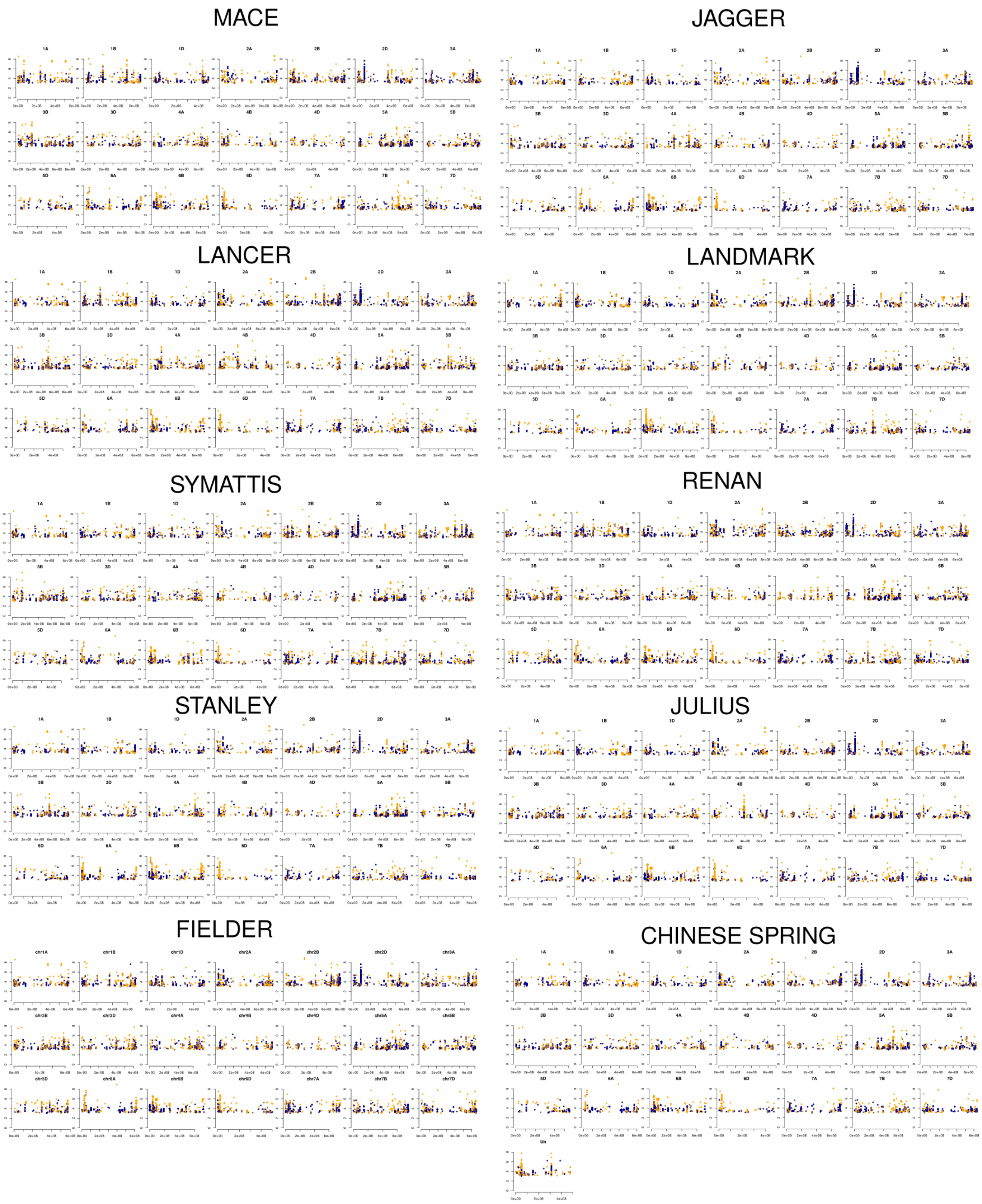


**Supplemental Figure 15**: Manhattan plot for field resistance to powdery mildew for the winter wheat field experiment 2023. The 10 reference genomes used in this study are represented. The colors represent the allele frequency of the *k*-mers with orange for extreme frequencies (close to 0 or 1) and blue for frequencies around 0.5

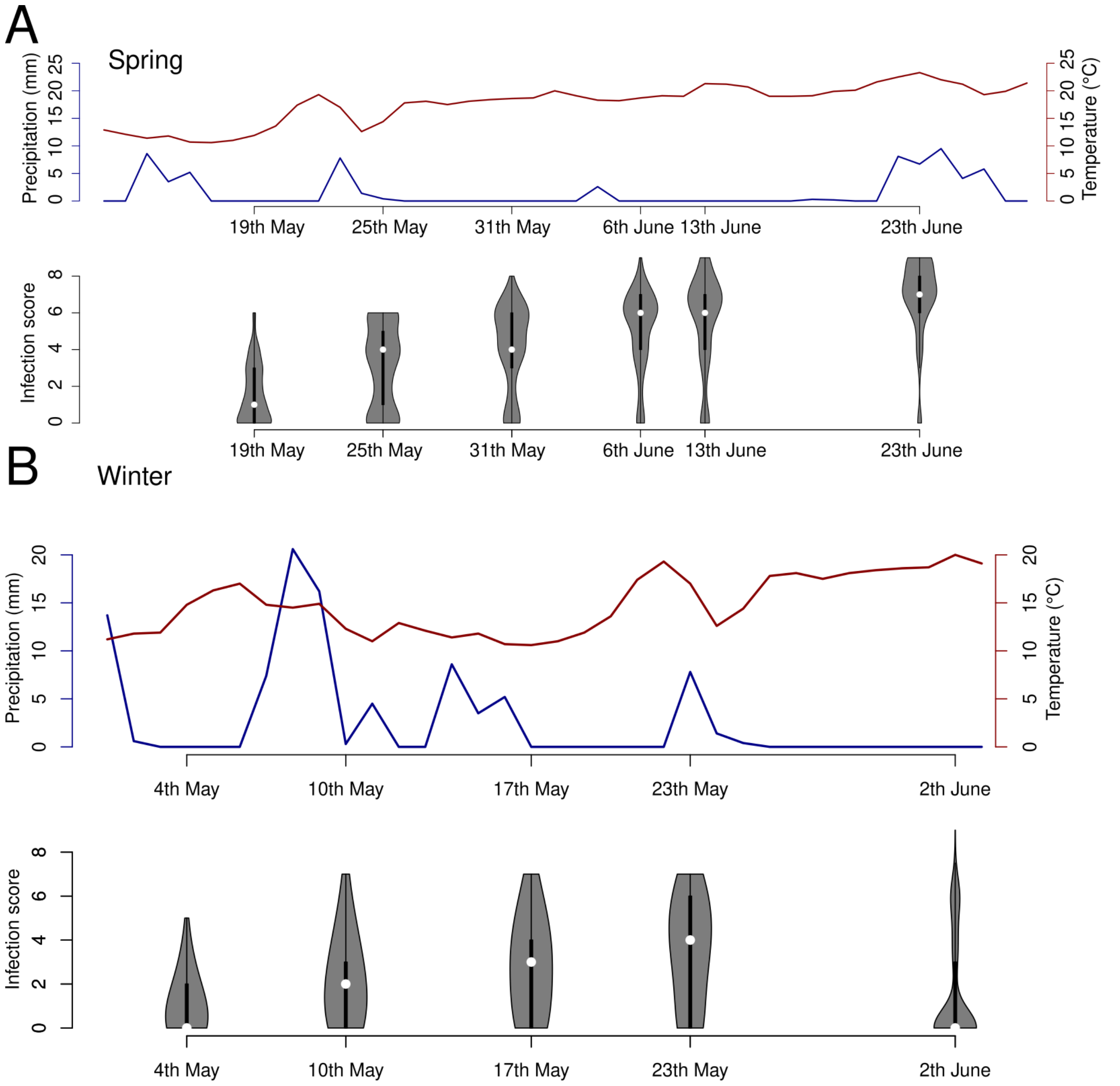


**Supplemental Figure 16**: Phenotype distribution of the field data 2023 separated based on the phenotyping dates with the corresponding temperature and precipitation for Spring (A) and Winter (B) wheat trial

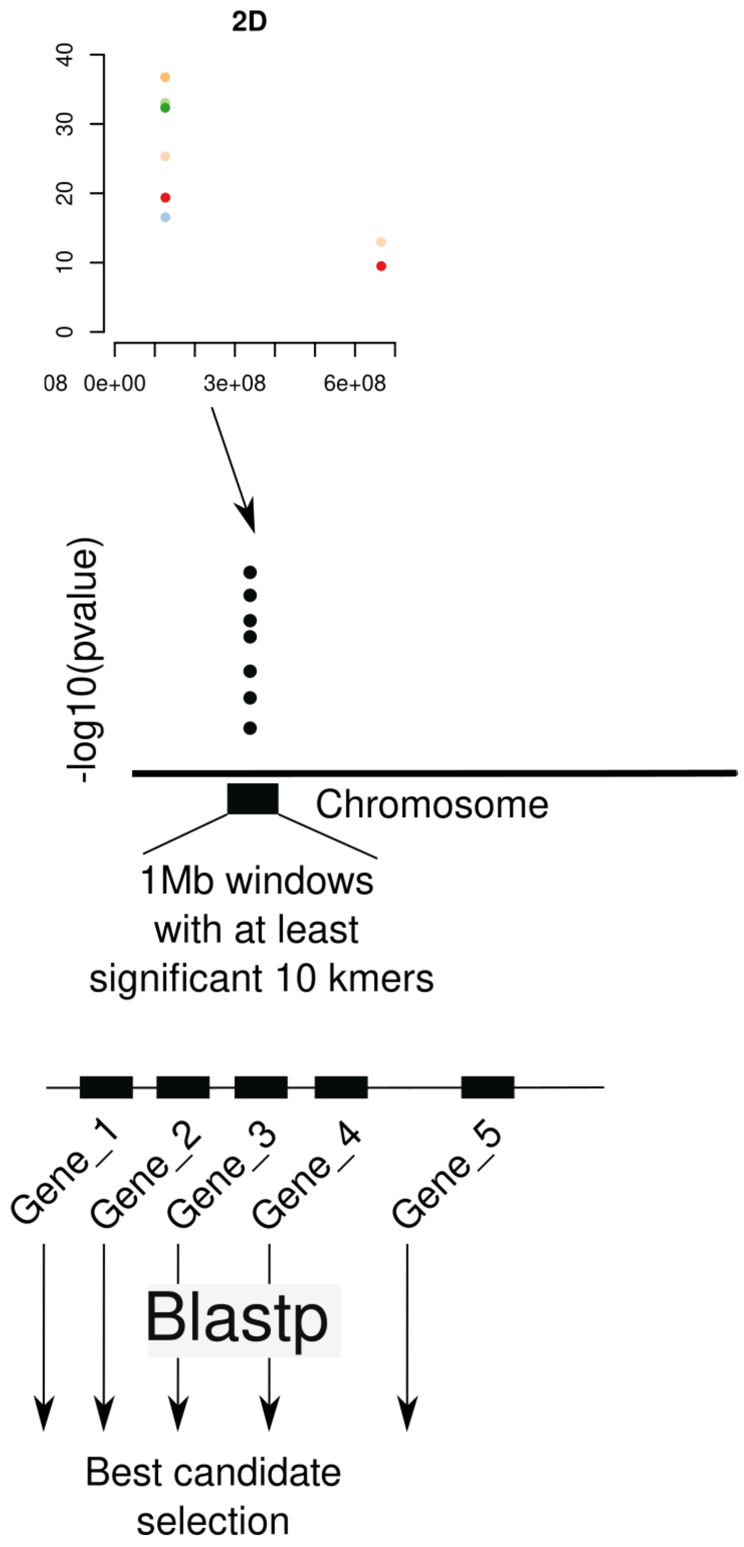


**Supplemental Figure 17**: Candidate selection approach: For each of the associated regions detected, a region of 1 Mb around the most significantly associated SNPs was extracted and all the protein sequences of the annotated genes within the region were extracted. The protein sequences of interest were then blasted to the NCBI database to assign a function to each gene (see Methods).

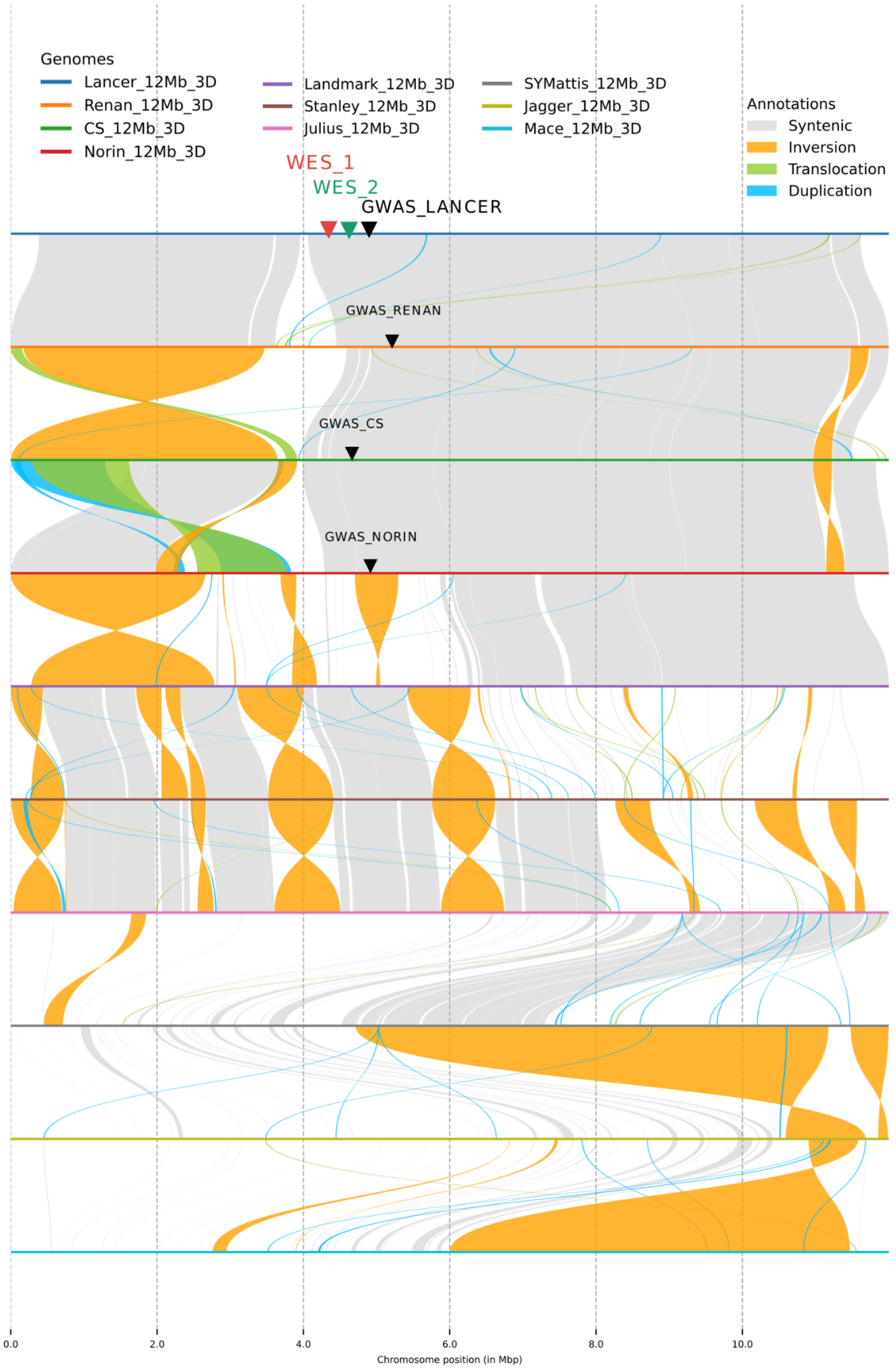


**Supplemental Figure 18**: Genome alignment of the first 12Mb of chromosome 3D for all the genomes. WES 1 and 2 represent the position of the Wax ester synthase candidate. Each section represents 2 Mb.
